# Dynamics and interplay of gene expression and chromosome organization across a predatory lifecycle

**DOI:** 10.1101/2025.09.24.678257

**Authors:** Renske van Raaphorst, Fares Osam Yáñez-Cuna, Corina Pascal, Ngat Tran, Jovana Kaljević, Swapnesh Panigrahi, Tung B. K. Le, Romain Koszul, Géraldine Laloux

**Affiliations:** de Duve Institute, UCLouvain, Brussels, Belgium; Institut Pasteur, CNRS UMR 3525, Université Paris Cité, Unité Régulation Spatiale des Génomes, Paris, France; John Innes Centre, Department of Molecular Microbiology, Norwich, United Kingdom; Laboratoire de Chimie Bactérienne, CNRS Marseille, France; WEL Research Institute, Wavre, Belgium; Sorbonne Université, Collège Doctoral, Paris, France

**Keywords:** Bacterial cell cycle, Chromosome organization, Bacterial predation, *Bdellovibrio bacteriovorus*, RNA-seq, Hi-C, ChIP-seq, Nucleoid, SMC condensin, ParABS

## Abstract

The obligate predatory bacterium *Bdellovibrio bacteriovorus* alternates between a motile attack phase and a growth phase inside another bacterium, during which it undergoes multiple rounds of DNA replication followed by non-binary division and progeny release. How its chromosome is dynamically reorganized across these contrasting states remains unclear. Here, we used synchronized predatory infections combined with time-resolved RNA-seq, Hi-C, ChIP-seq, and quantitative microscopy to map transcriptional activity and chromosome architecture across seven key stages of the predatory lifecycle. We uncover a dramatic shift in nucleoid organization – from an ultra-compact, transcriptionally restricted state in attack-phase cells to a progressively decompacted, *ori*-centered conformation during intracellular replication. Notably, distinct transcriptional and chromosome folding states not only mark the canonical attack and growth phases, but also critical transitions such as prey invasion, replication onset, and predator division. Temporal gene expression profiles reflect the coordinated regulation of core cellular functions across the predatory cycle. Furthermore, spatial localization of RNA polymerase, ribosomes, DNA-binding proteins, and freely diffusing proteins highlight the tight, cell cycle-dependent coupling between nucleoid compaction, chromosome accessibility, and transcriptional activity. Altogether, our findings reveal the intricate dynamics and coordination of genome folding and gene expression in a complex bacterial cell cycle driven by prey-predator interactions.

## INTRODUCTION

Optimal bacterial cell cycle progression and adaptation to fluctuating environments entail tight temporal control of gene expression. In parallel, chromosome organization plays a fundamental role in structuring the intracellular space and undergoes dynamic remodeling during key cellular events such as DNA replication, segregation, and differentiation^1,2^. How transcription and chromosome architecture are coordinated throughout the cell cycle remains largely unexplored ^3^. The predatory bacterium *Bdellovibrio bacteriovorus* provides a compelling model to investigate this interplay due to its complex yet synchronizable cell cycle, alternating between distinct non-replicative and replicative stages.

*B. bacteriovorus* is an obligate endobiotic predator of many diderm bacteria ^4^. Its lifecycle is typically divided into two main phases defined by the predation stage ^5–7^ (**Figure 1A**). During the non-proliferative attack phase (AP), it is a fast-swimming, non-replicative monoploid bacterium. Upon encountering a prey bacterium, *B. bacteriovorus* invades its periplasm, forming a specialized intracellular niche termed the bdelloplast ^8^. Here, the predator transitions into a growth phase (GP), elongating into a filament while consuming prey cellular contents, and ultimately dividing into a variable number of progeny proportional to prey cell size ^9,10^. Although the prey bacterium is killed early upon contact – and therefore not strictly a “host” –, the fact that *B. bacteriovorus* cell cycle progression depends on predation is reminiscent of certain host-pathogen interactions in which the pathogen cell cycle is tightly coupled to host invasion ^11–13^. How *B. bacteriovorus* orchestrates the switch between AP and GP states, coordinating these striking cell cycle transitions with prey invasion and digestion, remains poorly understood ^4,5^. Earlier transcriptomic analyses have shown distinct gene expression profiles between AP and GP, with fewer genes active in AP ^14^. We previously observed that the nucleoid in AP cells is tightly packed, partially excluding freely diffusing cytoplasmic proteins, suggesting restricted chromosome accessibility ^15^. Yet, it is unclear how the nucleoid transitions from a quiescent, highly compact state to a replicative state – and how chromosome organization is coordinated with gene expression throughout the predatory cycle.

**Figure 1.**
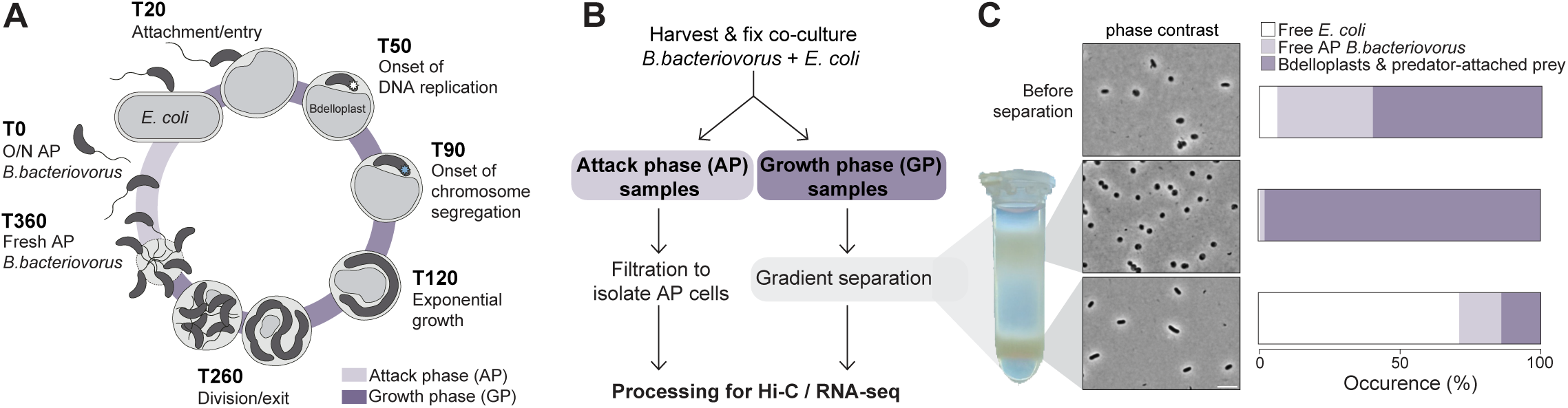
Synchronization of the predatory lifecycle of *B. bacteriovorus* for time-resolved transcriptomic and chromosome conformation analyses. **A.** Schematic representation of the *B. bacteriovorus* cell cycle and the time points sampled in this study. White star at T50 represents the first replisome; blue star at T90 represents the ParB·*parS* partitioning complex near the chromosomal origin. Prey rounding is typically observed shortly before entry of the predator, as a result of peptidoglycan modification. **B.** Experimental setup for Hi-C and RNA-seq sampling. **C**. Gradient separation 20 minutes after incubation leads to a sample (top band) containing 98% bdelloplasts and rounded *E. coli* with surface-attached predator. Counts for all time points are shown in Figure S1C. Representative experiment (repeated twice). **A-C.** The light and dark purple shades correspond respectively to free non-replicative predators (attack phase; AP samples), and predator-attached rounded prey and bdelloplasts (*B.* bacteriovorus-infected prey) collectively named growth phase (GP) samples.

The *B. bacteriovorus* chromosome exhibits a longitudinal *ori-ter* organization, with the origin of replication (*ori*) positioned near the prey-invasive cell pole ^15^. During the GP, a first round of replication and ParAB*s*-dependent segregation of duplicated *ori* is followed by asynchronous initiation of additional replication rounds, coupled with progressive segregation of sister chromosomes along the growing predator cell ^15–17^. This pattern yields odd or even chromosome copy numbers, regularly distributed in the filamentous mother cell before division ^15,16^. Apart from the role of the ParAB*s* system, active only in GP to partition *ori* copies ^16^, the broader organization of the chromosome throughout this cell cycle remains unknown. *B. bacteriovorus* encodes various homologs of nucleoid-associated proteins (NAPs), including the condensin SMC, two HU homologs (HUαβ), Fis, IHFαβ, two Dps homologs, and the recently identified histone-fold proteins Bd0055 and Bd3044 ^18,19^. While Bd0055 is essential for survival, its *in vivo* function is unknown ^18,19^, and all other NAPs remain unexplored.

In model bacteria like *E. coli*, chromosome folding is profoundly affected by gene expression ^20^. Highly transcribed loci, such as ribosomal RNA (rRNA) operons, typically localize at the nucleoid periphery during exponential growth ^21,22^, and inhibitors of transcription or translation dramatically alter nucleoid compaction ^23–26^. High-resolution chromosome contact maps obtained through Hi-C experiments have demonstrated that transcription imposes local constraints shaping three-dimensional chromosome conformation at larger scales ^27,28^. Additionally, NAPs can influence this transcription-related organization ^29–31^. Although previous transcriptomic studies have investigated AP and GP stages in *B. bacteriovorus* ^14,32^, a time-resolved picture of gene expression spanning the entire predatory lifecycle is still lacking.

Here we filled these gaps by employing an integrative approach combining RNA-sequencing (RNA-seq), high-resolution chromosome conformation capture (Hi-C), chromatin immunoprecipitation followed by sequencing (ChIP-seq), and quantitative microscopy analyses to simultaneously examine chromosome organization and transcription at seven key steps of the synchronized *B. bacteriovorus* lifecycle. Our results provide insights into the dynamic interplay between nucleoid conformation and gene expression, uncovering cell cycle transitions beyond the traditional biphasic predator-prey interaction model.

## RESULTS

### Synchronization of predation for time-resolved transcriptomics and chromosome conformation capture

To capture the full *B. bacteriovorus* lifecycle, we selected seven time points covering key events (T, in minutes; **Figure 1A**): attack phase (AP) from an overnight culture (T0), prey attachment and entry (T20), initiation of DNA replication (T50), onset of chromosome segregation (T90), mid-growth (T120), predator division and exit (T260), and newly released AP cells (T360). Replication and segregation timing were guided by previous work ^15,16^ and verified using fluorescent fusions to DnaN and ParB ^15,16^ (**Figure S1A-B**).

Synchronization of the *B. bacteriovorus* predatory cycle is possible because prey invasion by fresh AP predators occurs rapidly and in a narrow time window upon mixing with *Escherichia coli*. Using an excess of prey further ensures near-simultaneous prey invasion by the entire predator population ^9,15,16^. To isolate synchronized predator populations throughout the lifecycle, we leveraged the differential migration of “infected” and “uninfected” *E. coli* in a density gradient (**Figure 1B-C**). This allowed us to recover bdelloplasts – and prey with surface-attached predators at the earliest stage – at the selected time points (T20-T260) following predator-prey mixing at a 1:2 ratio, while free AP predators (T0 and T360) were isolated by size filtration as previously described ^33^ (**Figure 1B-C**). Phase contrast microscopy confirmed that non-AP samples typically contained >95% bdelloplasts (**Figure 1C, Figure S1C**).

All collected samples underwent RNA-seq and Hi-C analyses. Mapping reads from both analyses to the *E. coli* and *B. bacteriovorus* genomes confirmed robust synchronization of the predatory cycle (**Figures S2A, S3A**). Early time points (T20 to T90) contained a higher proportion of *E. coli* reads, reflecting abundance and high DNA content of prey cultured under fast growth conditions before predation, whereas later stages showed fewer or negligible *E. coli* reads due to progressive digestion ^34^ (**Figures S2A, S3A**). Conversely, the fraction of reads mapping to *B. bacteriovorus* increased throughout the GP time points, consistent with predator proliferation.

### Gene expression exhibits global changes during the predatory lifecycle

Replicates from non-proliferative phases (T0 and T360) strongly correlate in both their RNA-seq profiles (**Figure S2B-C**, Pearson R > 0.5) and Hi-C datasets (**Figure S3B**, Pearson R > 0.98, assessed by HiCRep ^35^), indicating that these timepoints correspond to similar physiological states and consistent with our sampling of a complete predation cycle. Conversely, time points corresponding to prey invasion and GP differ substantially, especially when comparing gene expression profiles at initial invasion (T20) and late GP (T260) stages (Pearson R < 0.1, **Figure S2B-C**). Notably, a global burst of gene expression occurred upon prey attachment and entry (T20), whereas extensive transcriptional shutdown characterized the division and exit phase (T260) (**Figure 2A-B**). Thus, the global gene expression landscape of *B. bacteriovorus* fluctuates extensively during its predatory lifecycle, with the observed dynamic profiles clearly reflecting three major phases in the predatory lifecycle: prey attachment and entry, predator growth and replication, and predator division and escape from prey.

**Figure 2.**
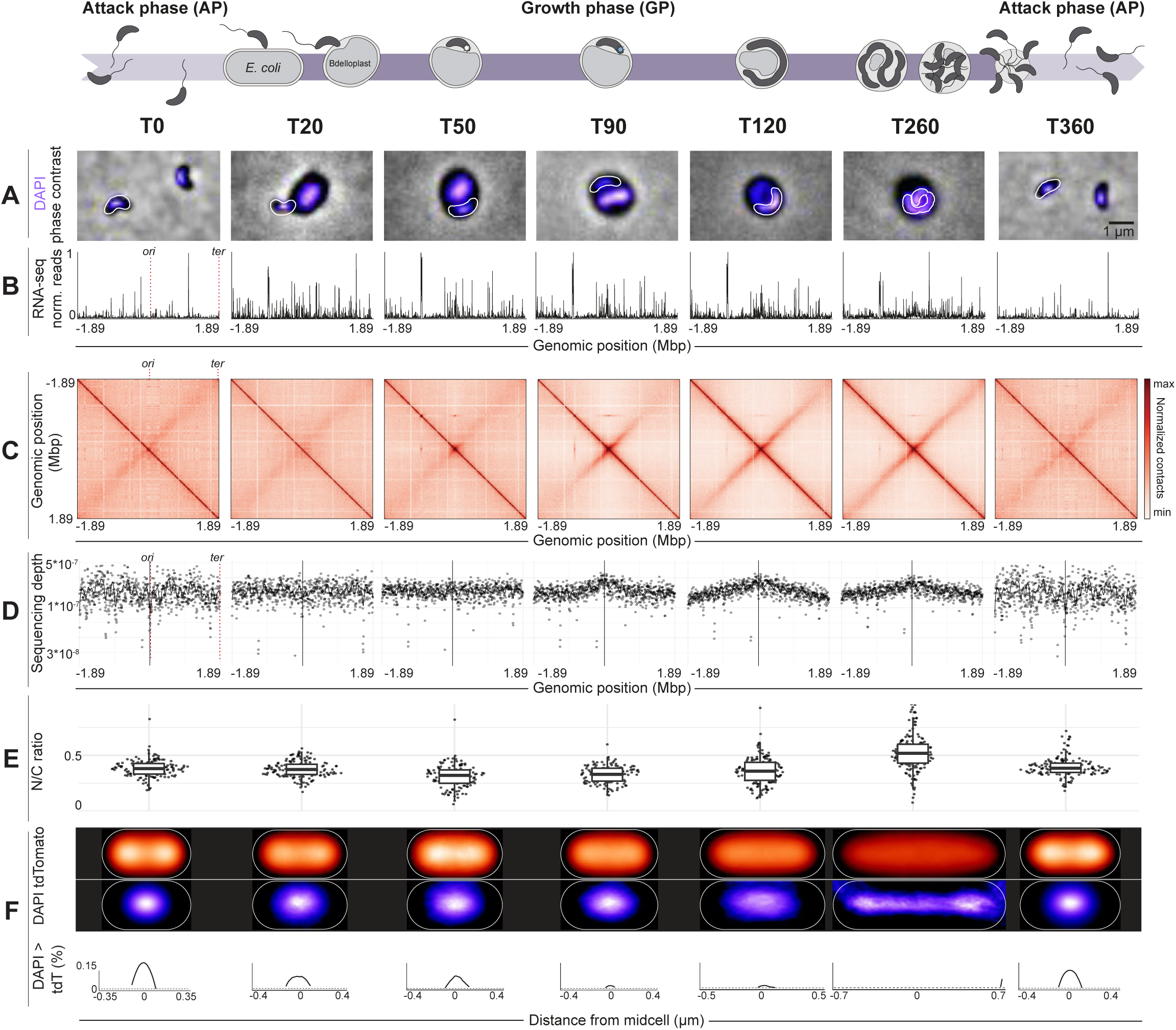
Global changes in nucleoid conformation, topology of gene expression, and nucleocytoplasmic ratio during the *B. bacteriovorus* lifecycle. Top: schematics of *B. bacteriovorus* cell cycle progression on top of the corresponding sampled time points; white and blue stars represent the first replisome and ParB·*parS* partitioning complex as in Figure 1A. **A.** Representative microscopy images of *B. bacteriovorus* infecting *E. coli* cells 0, 20, 50, 90, 120, 260 and 360 minutes after co-incubation; composite image (purple: DAPI, grey: phase contrast). **B.** Global gene expression landscape: distribution of normalized (0-1) RNA-seq reads on the circular chromosome, where the *ori* is positioned at 0 Mbp. **C.** Chromosome conformation maps derived from Hi-C of each corresponding time point, centered on the *ori* (position 0 Mbp); color intensity scales with contact frequency. **D.** Sequencing coverage along the *B. bacteriovorus* chromosome at each corresponding time point, with T50 representing the onset of DNA replication initiation with a small peak at the *ori*, which is positioned at the center of the graph as indicated by the vertical line; coverage data were derived from the Hi-C sequencing data. **E.** The nucleocytoplasmic (N/C) ratio at each time point shows a relative increase in nucleoid to cell size ratio (using measurements of respective areas in individual predator cells) from T120 to T260. **F.** Top: median distribution of fluorescent signals from free cytosolic tdTomato (red) and DNA (stained with DAPI, purple) in *B. bacteriovorus* cells (strain GL1462), measured by microscopy at each time point, shows gradual loss of nucleoid exclusion of cytosolic proteins (seen as bilobed distribution of the tdTomato signal), starting from T20 (attachment/entry). Median distributions of each time point were obtained by straightening the cell intensity profiles and calculating the median fluorescence intensity per relative length/width localization in the cell. Bottom: distribution difference where DAPI > tdTomato, calculated by subtracting the median fluorescence intensities of DAPI and tdTomato over the length axis of the cell, and plotting the points where DAPI > tdTomato.

### Chromosome conformation shifts during predation and cell cycle progression

Concurrent Hi-C datasets revealed striking chromosome rearrangements across the lifecycle. Non-replicative stages (T0, T20, and T360) exhibited a highly compacted nucleoid, shown as homogenous, densely red Hi-C maps with a thin main diagonal (**Figure 2C-D**, **Figure S4**). These maps are dominated by extensive and non-specific long-range contacts, consistent with a strong compaction of the chromosome ^15^. In contrast, replicative stages (T50-T260) were characterized by nucleoid decompaction. This is highlighted by enriched short- and medium-range contacts near the *ori*, accompanied by reduced long-range contacts and broadening of the main diagonal, progressively extending towards the terminus (*ter*) until T120 (**Figure 2C-D**, **Figure S4**). Furthermore, a clear inter-arm interaction emerged from the *ori*, perpendicular to the main diagonal (T50; **Figure 2C, Figure S4**) – a pattern typically indicating ParB-mediated recruitment of SMC proteins to the *ori*, bridging the two replichores during replication ^36–39^. Consistent with this model, expression of the *B. bacteriovorus* SMC homolog peaked during replication stages (see below), and the secondary diagonal coincided with the timing of ParB localization at the *ori* region (^16^ and **Figure S1A**). Interestingly, a faint inter-arm interaction persisted even in non-replicating cells (T0, T20, and T360; **Figure 2C-D, Figure S4**). This suggests that despite nucleoid compaction and absence of ParB and SMC proteins at that stage ^16,18^, the non-replicating chromosome maintains a longitudinal organization, in agreement with the observed *ori* and *ter* positions in AP cells ^15^. Collectively, our findings demonstrate that the three-dimensional architecture of the *B. bacteriovorus* chromosome shifts from highly compacted during the AP to increasingly decompacted upon initiation and progression of replication within prey cells.

### Nucleoid expansion and accessibility increase during the growth phase within prey

To monitor changes in chromosome organization across the *B. bacteriovorus* lifecycle at the single-cell level, we quantified the nucleocytoplasmic (N/C) ratio, defined as the fraction of predator cell area occupied by DNA. Automated measurement of predator cell dimensions within prey is challenging when using classical image segmentation tools ^9^; thus, we trained the deep learning segmentation network MiSiC ^40^ to recognize predator cells based on diffuse cytoplasmic fluorescence (**Figure S5**). Additionally, we denoised the DAPI images using the Noise2Void deep learning network ^41^ to overcome interference from the stronger, neighboring *E. coli* nucleoid signal ^32^ (**Figure S5**). Applying this approach to DAPI-stained *B. bacteriovorus* cells constitutively producing cytosolic tdTomato ^9^, we observed nucleoid expansion (increased N/C ratio) only after chromosome segregation had initiated (**Figure 2E**, T120-T260). To determine whether nucleoid expansion correlates with increased accessibility, we compared the spatial distributions of cytoplasmic tdTomato and DAPI signals in predator cells (**Figure 2F**). Nucleoid exclusion of cytosolic proteins, assessed by the difference between these two reporters, was most pronounced in the AP samples (T0 and T360) but decreased when predators invade prey (**Figure 2F**), before DNA replication (**Figure 2D**) and visible chromosome expansion (**Figure 2C,E**). This aligns with previous observations using free fluorescent proteins ^15^. The NAP HUα, tagged with msfGFP and used as a marker for nucleoid localization, displayed a spatial distribution similar to DAPI, ruling out possible nucleoid condensation artefacts from the staining (**Figure S6**). These data indicate that chromosome accessibility to freely diffusing proteins begins to increase prior to visible nucleoid expansion during intraperiplasmic predator growth.

### Transcription correlates with chromosome compaction exclusively during the growth phase

Transcription commonly influences chromosome folding locally, with positive correlations observed between transcriptional activity and enriched contacts among adjacent loci in several bacterial species ^27,31^. We found that in *B. bacteriovorus*, this relationship changes throughout the cell cycle. During the AP, when transcriptional activity is globally reduced and the compact chromosome presents a relatively uniform contact pattern (**Figure 2B-C**), there was no detectable correlation (T0; Pearson R = 0.05; **Figure S4**), as expected between signals with minimal fluctuations. However, upon prey invasion, when transcription markedly increases (**Figure 2B**), a correlation with chromosome contacts became apparent (T20; Pearson R = 0.28), reaching a peak at the onset of DNA replication (T50; Pearson R = 0.50) and gradually decreasing thereafter (**Figure S4**). Strikingly, intense hubs of short-range contacts emerged at T20 and T50 around the *ori* and at a locus approximately three-quarters along the chromosome, which contains several rRNA operons (**Figure S4**, arrows). These short-range contact hubs coincided with a burst of transcriptional activity at these loci, and persisted during growth (**Figure 3F**, arrows; **Figure S4**). In addition, Hi-C maps revealed that these highly expressed loci also formed a discrete long-range interaction spanning a ∼900 kb region (**Figure 3F**, arrows; **Figure S4**). This spatial configuration is reminiscent of the colocalization of rRNA operons in *E. coli* during exponential growth – which depends entirely on their extremely high expression level ^14^ – supporting the idea that active genes associate in space, potentially due to their repositioning in regions of high transcriptional activity at the nucleoid periphery ^27^.

**Figure 3.**
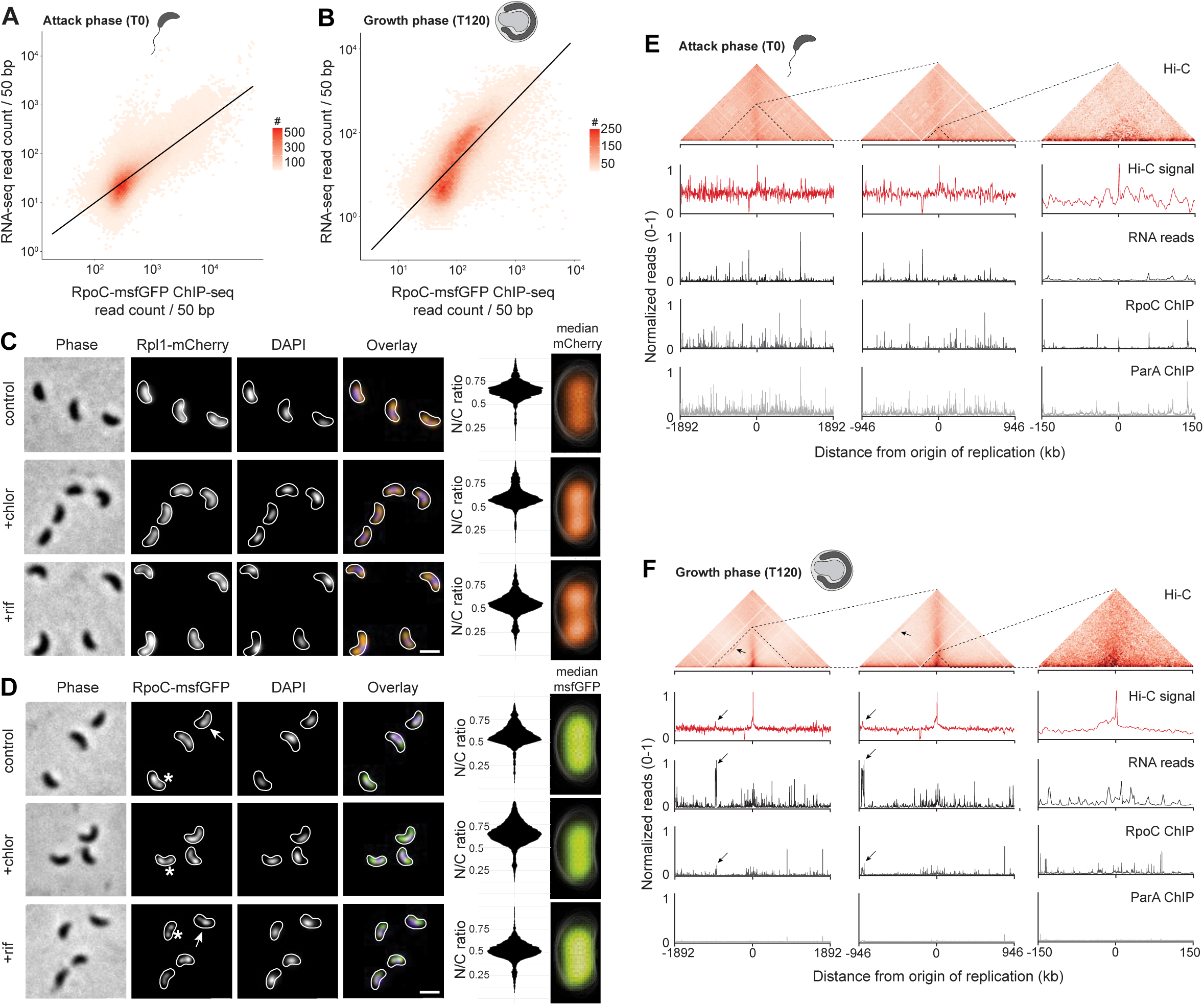
Time-dependent comparisons between RNA-seq reads, RNA polymerase and ParA localization, and chromosome contacts. **A-B**. RpoC-msfGFP ChIP-seq correlation with RNA-seq in AP (A) and during growth (T120, B). Read counts of RpoC-msfGFP ChIP-seq and RNA-seq were binned per 50 base pairs. Black line = linear model; Pearson R in AP (A, T0) = 0.62, Pearson R in GP (B, T120) = 0.52. **C-D.** Localization of Rpl1-mCherry (labeling the L1 ribosomal protein; strain GL1027) (C), RpoC-msfGFP (labeling the RNA polymerase; strain GL1026) (D), and nucleoid size do not change upon addition of chloramphenicol or rifampicin. Representative images, calculated N/C ratio, and median fluorescence signal of attack phase cells treated with 600 µg/mL chloramphenicol or 150 µg/mL rifampicin, respectively, for 4 hours. Stars: nucleoid-excluded RpoC-msfGFP. Arrows: nucleoid-localized RpoC-msfGFP. Scale bar = 1 µm. **E-F.** Pile-up of (top to bottom): Hi-C contact map, normalized Hi-C signal (red), RNA-seq reads (black), RpoC-msfGFP ChIP-seq reads (IP/input per 500 base pairs; dark grey) and ParA-msfGFP ChIP-seq reads (IP/input per 500 base pairs; light grey), in AP (T0, E) and GP (T120, F). Left to right zooms in from the full genome (left) to –946:946 kb distance from *ori* (middle) to –150:150 kb distance from *ori* (right). Arrows point to the correlated gene expression burst and increased contacts at the three-quarter position of the chromosome.

### Transcription is spatially restricted to accessible chromosomal loci in non-replicating predators

Given the limited accessibility of the nucleoid to freely diffusing proteins during the AP (^15^ and **Figure 2**), we next asked how transcription and translation are spatially arranged in *B. bacteriovorus*. Because RNA abundance may not directly reflect active transcription due to effects of RNA stability, we assessed chromosomal occupancy by the RNA polymerase via chromatin immunoprecipitation followed by sequencing (ChIP-seq) on cells producing the RNA polymerase subunit C (RpoC) fused to msfGFP during AP (T0) and GP (T120). The positive correlation between ChIP-seq and RNA-seq reads during these two stages of the cell cycle (Pearson R = 0.52 (T0) and 0.62 (T120); **Figure 3A-B**) indicates that RNA-seq largely captures loci of active transcription. To further assess the subcellular organization of transcription and translation, we localized natively produced fluorescent fusions to RpoC and the ribosomal protein L1 (Rpl1). Rpl1-mCherry was partially excluded from the nucleoid (**Figure 3C**), reminiscent of patterns observed for freely diffusing cytosolic proteins. In contrast, RpoC-msfGFP accumulated on and around the nucleoid (**Figure 3D**, arrows), occasionally exhibiting mid-cell exclusion patterns (**Figure 3D**, stars) similar to the non-specific DNA-binding protein ParA during AP ^16^. Consistent with this observation, the ParA ChIP-seq profile closely matched that of RpoC at T0 (**Figures 3E**) and correlated with the RNA-seq profile at this cell cycle stage (ParA-msfGFP ChIP-seq vs RNA-seq at T0, Pearson R = 0.5; **Figure S7A**). This was not the case during the GP (ParA-msfGFP ChIP-seq vs RNA-seq at T120, Pearson R = 0.26; **Figures 3F, S7B**), as expected since ParA drives chromosome segregation in replicating cells via interactions with both the chromosome and *ori*-localized ParB ^16^. Together, these data suggest that during the non-proliferative phase of the *B. bacteriovorus* cell cycle, transcription predominantly occurs at regions of the compacted nucleoid that are accessible to non-specific DNA-binding proteins. Whether this pattern arises from transcription being constrained by local chromosome accessibility or from active transcription itself facilitating local accessibility remains to be fully elucidated.

### Transcription and translation are not the main drivers of nucleoid compaction in attack phase

To directly probe the link between nucleoid compaction, transcription, and translation, we inhibited these processes in newborn AP *B. bacteriovorus* cells using high concentrations of rifampicin (150 µg/mL) or chloramphenicol (600 µg/mL), respectively. Unlike observations in other bacteria ^26,42,43^, the N/C ratio in AP *B. bacteriovorus* cells did not change noticeably upon treatment with either antibiotic, nor did the subcellular localization of Rpl1 and RpoC (**Figure 3C-D**). AP cells remained motile for at least 4 hours post-treatment, and viability and predatory capacity were unchanged following 8-hour incubation with either drug, as measured by plaque-forming assays (control: 2 × 10^7^ PFU/mL; chloramphenicol: 0.9 × 10^7^ PFU/mL; rifampicin: 3 × 10^7^ PFU/mL) – consistent with overall low gene expression activities at that stage. As lower concentrations of these antibiotics halted cell cycle progression during GP (MIC rifampicin = 0.085 µg/mL, MIC chloramphenicol = 0.15 µg/mL; **Figure S8**), our results strongly suggest that transcription and translation are not the main drivers of nucleoid compaction in AP predator cells.

### Distinct gene expression clusters highlight cell cycle stages and transitions

Our time-resolved RNA-seq dataset allowed us to capture gene expression at defined stages across the predatory cycle. Hierarchical clustering of genes differentially expressed from T0 to T260 compared to newborn predator cells (T360) yielded 11 gene clusters, each representing a distinct temporal expression pattern reflecting particular functional roles (**Figure 4A**). For instance, genes in cluster 2 show peak expression at T260 – coinciding with cell division and prey exit – and are enriched in chemotaxis and flagellar assembly functions required for prey hunting ^44^, as well as cell wall organization and morphology (**Figure 4B**). By contrast, genes in cluster 8 peak early at prey invasion (T20) and predominantly encode components involved in transcription and translation, including several highly expressed rRNA operons (**Figure 4C**). Intriguingly, these genes are primarily located near the chromosomal *ori* and the three-quarter region that strongly interacts with the *ori* during growth (T20-T260; **Figure 2C, Figure S4**), likely supporting the early burst of protein synthesis required for predator growth, DNA replication, and prey digestion.

**Figure 4.**
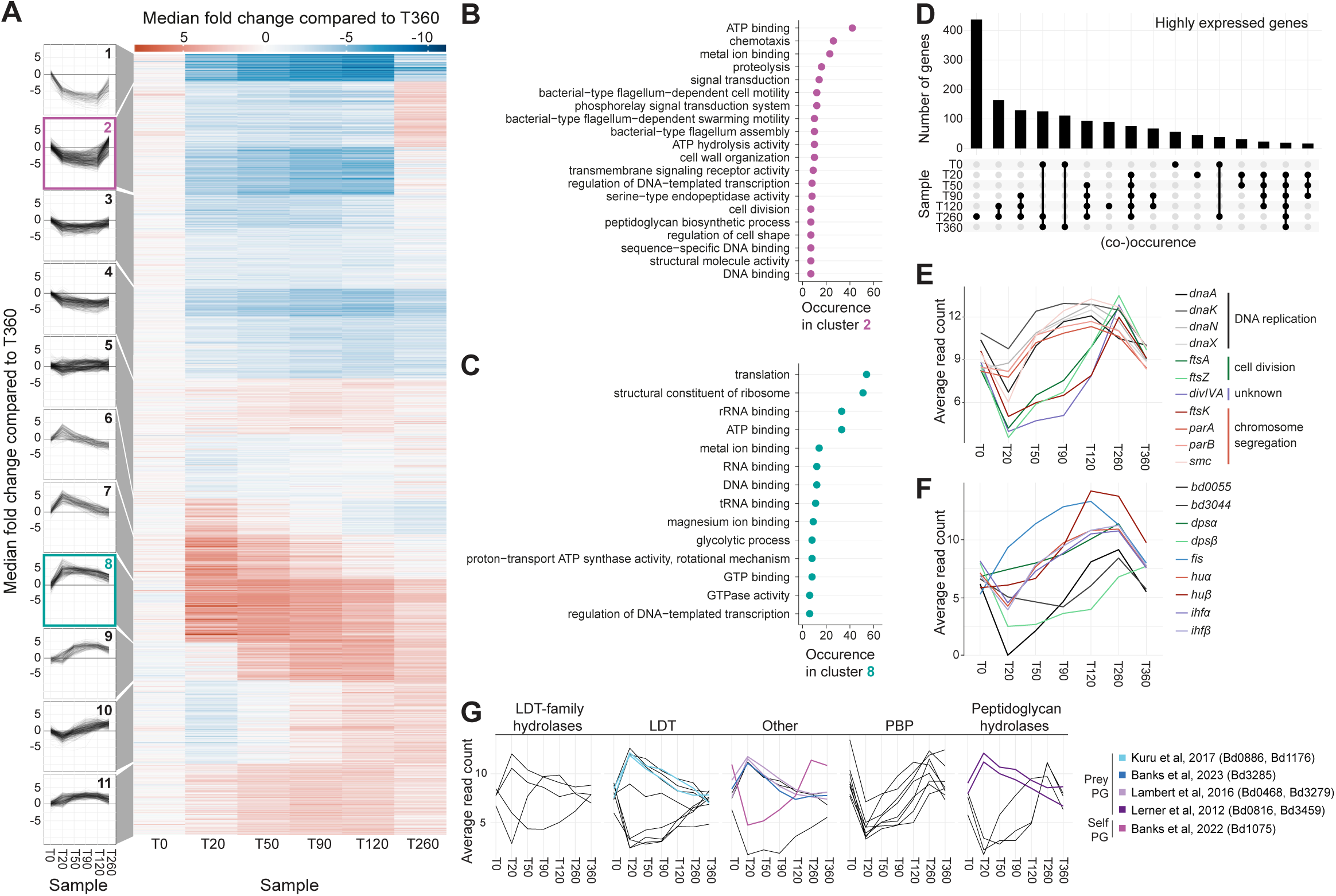
Gene expression profiles during the *B. bacteriovorus* predatory lifecycle. **A.** Clustering of differential gene expression profiles by hierarchical K-means shows 11 temporal expression profiles. Left: grouped expression profiles, with clusters 2 and 8 highlighted and further visualized in B and C. Right: fold change compared to T360 (fresh AP), grouped per cluster. **B.** Top Gene Ontology (GO) classes occurring in cluster 2. GO biological and molecular function ontologies occurring > 5 times. **C.** Top GO classes occurring in cluster 8. GO biological and molecular function ontologies occurring > 5 times. **D.** UpSet plot showing the number of genes with high expression (median counts > 10) at one or more specific time points. **E.** Normalized median expression profiles of genes involved in DNA replication, cell division, and chromosome segregation. Average read counts over time; colors indicate the associated cell cycle process. **F.** Normalized median expression of genes encoding nucleoid associated proteins (NAPs). **G.** Normalized median expression profiles of genes involved in peptidoglycan modification (average read counts over time). Genes are faceted by (predicted) function; colors indicate previous publications about said gene (see references in text). LDT, L,D-transpeptidases; PBP: penicillin-binding proteins.

Notably, Hocher et al. proposed that proteins present in AP predators are largely synthesized during the preceding growth phase ^18^. The AP proteome in this study indeed correlates best with our RNA-seq profile obtained at the last GP time point (**Figure S9**, T260).

Highly co-expressed genes often share related biological roles. The latest growth stage (T260) displayed the largest group of uniquely highly expressed genes (>400 genes, fold change ≥ 5, p-value ≤ 0.01; **Figure 4D**). These include genes encoding the key cell division protein FtsZ as well as DivIVA, shown to localize at constriction sites prior to predator cell division ^45^ (**Figure 4E**), along with genes encoding putative Type IV filaments proteins and flagellins. This highlights not only processes occurring only at the end of the GP (such as cell division) but also the preparation of future progenies with features important for the upcoming prey hunting stage ^46^.

Key cell cycle regulators exhibit precisely timed expression consistent with their functional roles (**Figure 4E**). For instance, genes required for DNA replication and chromosome segregation (*dnaA, dnaN, parA, parB, smc*) were strongly upregulated early in the GP, while those involved in late segregation and cell division (*ftsK, ftsZ, ftsA*) peaked late in the GP (T260). Likewise, expression profiles of NAPs showed distinct temporal patterns. Most predicted NAP genes were upregulated during growth and replication, matching the expression patterns of *parB* and *smc* (**Figure 4E-F**). However, genes encoding the histone-like proteins (*bd0055* and *bd3044*) ^18,19^ and the small, non-specific DNA-binding protein HUβ (*bd3383*) peaked specifically at T260 (**Figure 4F**), whereas DpsB (*bd2620*) is the only NAP that was upregulated during AP (T0 and T360, **Figure 4F**). These data indicate that while some NAPs may play redundant roles, they may have specialized roles in chromosome organization at defined stages or transitions of the *B. bacteriovorus* lifecycle.

### Self- vs prey-modifying enzymes display temporally distinct expression profiles

Examining precise gene expression profiles during the synchronized *B. bacteriovorus* lifecycle can provide clues on potential functions in key predation processes such as prey invasion, predator growth, or exit from prey. As a proof of concept, we compiled genes linked to peptidoglycan (PG) modification. Several enzymes previously implicated in prey PG remodeling showed peak gene expression specifically at T20, in line with their role in prey invasion **(Figure 4G)**. These included carboxypeptidases involved in prey rounding ^47^ (*bd3459* and *bd0816*), a lytic transglycosylase cleaving septal prey PG ^48^ (*bd3285*), L,D-transpeptidases resealing the predator entry site in the prey cell wall ^8^ (*bd1176* and *bd0886*), and N-acetylglucosamine deacetylases sensitizing prey PG for subsequent prey lysis at the end of the GP ^49,50^ (*bd3279* and *bd0468*). Conversely, Bd1075, implicated in predator cell curvature ^51^ rather than prey modification, is mainly expressed later in the GP (**Figure 4G**). Additionally, we identified two predicted L,D-transpeptidase family hydrolases (*bd0553*, *bd2061*) and three other L,D-transpeptidases (*bd3158*, *bd3176*, *bd3741*) whose peak expression at T20 suggests previously unrecognized roles in predatory processes.

## DISCUSSION

The lifecycle of *B. bacteriovorus* has traditionally been viewed as biphasic based on its relationship to the prey – an attack phase outside the prey and a growth phase inside it. Our study refines this view by revealing that *B. bacteriovorus* undergoes dramatic shifts in chromosome organization as it transitions between these two extremes, reflecting distinct nucleoid configurations and transcriptional profiles likely suited to nutrient-limited versus intracellular conditions. Similar genome reorganization may exist in other bacteria alternating between free-living and host-associated lifestyles. Importantly, our time-resolved analyses uncover unique gene expression and chromosome organization signatures during critical transitions, such as prey attachment, invasion, predator growth, division, and exit, extending beyond a simple biphasic model. Gene expression peaks identified in our study precede or coincide with recently reported protein abundance fluctuations during the *B. bacteriovorus* lifecycle ^52^, consistent with the expected delay between these processes. The detailed temporal map of the predatory cycle identified here therefore provides a powerful resource for dissecting the genetic regulation underlying predator-prey interactions and predator proliferation.

Previous bacterial Hi-C studies have mostly provided snapshots of chromosomal architecture in unsynchronized populations or at a given growth or developmental stage ^36,38,53–55^. Our Hi-C maps on synchronized populations provide an unprecedented step-by-step view of chromosome conformation dynamics across an entire bacterial cell cycle. Despite asynchronous replication cycles ^15^ and variable progeny numbers ^9^ in *B. bacteriovorus* (two to eight under our conditions, average = 2.32 ± 1.32), clear temporal patterns of global and local chromosome organization emerged. As the bacterium transitions from AP to intracellular growth and replication, we observed a remarkable shift from a uniformly dense nucleoid structure to an *ori*-centered conformation featuring inter-arm bridging conformation. In addition, prominent long-range contacts between highly expressed rRNA gene loci and the *ori* throughout the GP highlight intimate connections between transcriptional activity and chromosome topology.

An intriguing feature of the *B. bacteriovorus* lifecycle is the exceptionally compact nucleoid state in non-replicating cells, which suggests dedicated cell cycle-dependent mechanisms of chromosomal organization. Our RNA-seq analysis highlights a subset of NAPs as potential contributors to this unusual degree of compaction. For instance, several NAP genes were highly expressed during the late growth phase, suggesting that they may function in preparing *B. bacteriovorus* for its subsequent attack phase by establishing a quiescent nucleoid architecture. Among them, the essential bacterial histone Bd0055 does not compact the *E. coli* chromosome ^18^, possibly indicating species-specific properties or additional factors. Future *in vivo* investigation of the full set of NAPs in *B. bacteriovorus* may elucidate the underpinnings of its dynamic compaction state during the cell cycle.

Our findings also challenge existing models regarding nucleoid compaction determinants. Unlike *E. coli*, where transcription and translation profoundly affect nucleoid structure ^27,56^, our experiments with inhibitors hint that chromosome compaction in AP *B. bacteriovorus* cells is largely independent of active transcription and translation. This is consistent with the absence of correlation between short-range chromosome contacts and the RNA-seq profile, and the minimal transcriptional output observed at this stage (in agreement with a previous transcriptomic study ^14^). Instead, we propose that nucleoid compaction contributes to the global expression shutdown in AP, serving as a resource-saving adaptation during these nutrient limitation conditions ^57,58^, thereby conserving energy for the costly flagellar motility until prey encounter.

Finally, our integration of deep learning-based tools allowed us to measure changes in predator and nucleoid morphology at the single-cell level, significantly improving quantitative image analysis of filamentous intracellular *B. bacteriovorus*. This approach will also facilitate automated analysis of subcellular protein localization, which previously required manual outlining of the predator cell ^15,16,45^. More broadly, these tools will be valuable for probing the cell biology of other endobiotic predators or intracellular pathogens in the future.

In conclusion, our study reveals new facets of bacterial proliferation through a comprehensive, time-resolved exploration of transcription, nucleoid architecture, and cell cycle progression across a predatory lifecycle. The intricate connections between these processes not only illuminate features unique to the predatory bacteria studied here but also offer broader insights into bacterial chromosome biology. Our findings set the stage for investigating the molecular mechanisms of predator-prey interactions and chromosome organization in diverse bacterial species.

## MATERIALS & METHODS

### Growth media & culture

The strains used in this study are listed below. *E. coli* was routinely grown in Lysogeny Broth (LB, MP Biomedicals) at 37°C with aeration. Plasmids were maintained by adding kanamycin (50 μg/mL)*. B. bacteriovorus* strains were routinely grown in DNB medium (Dilute Nutrient Broth, Becton, Dickinson and Company), supplemented with 2 mM CaCl_2_ and 3 mM MgCl_2_ salts with *E. coli* MG1655 as prey at 30°C and constant shaking as previously described ^33^. When selecting *B. bacteriovorus* for kanamycin resistance, an MG1655 strain carrying a plasmid with a KanR cassette (GL818) was used as prey instead.

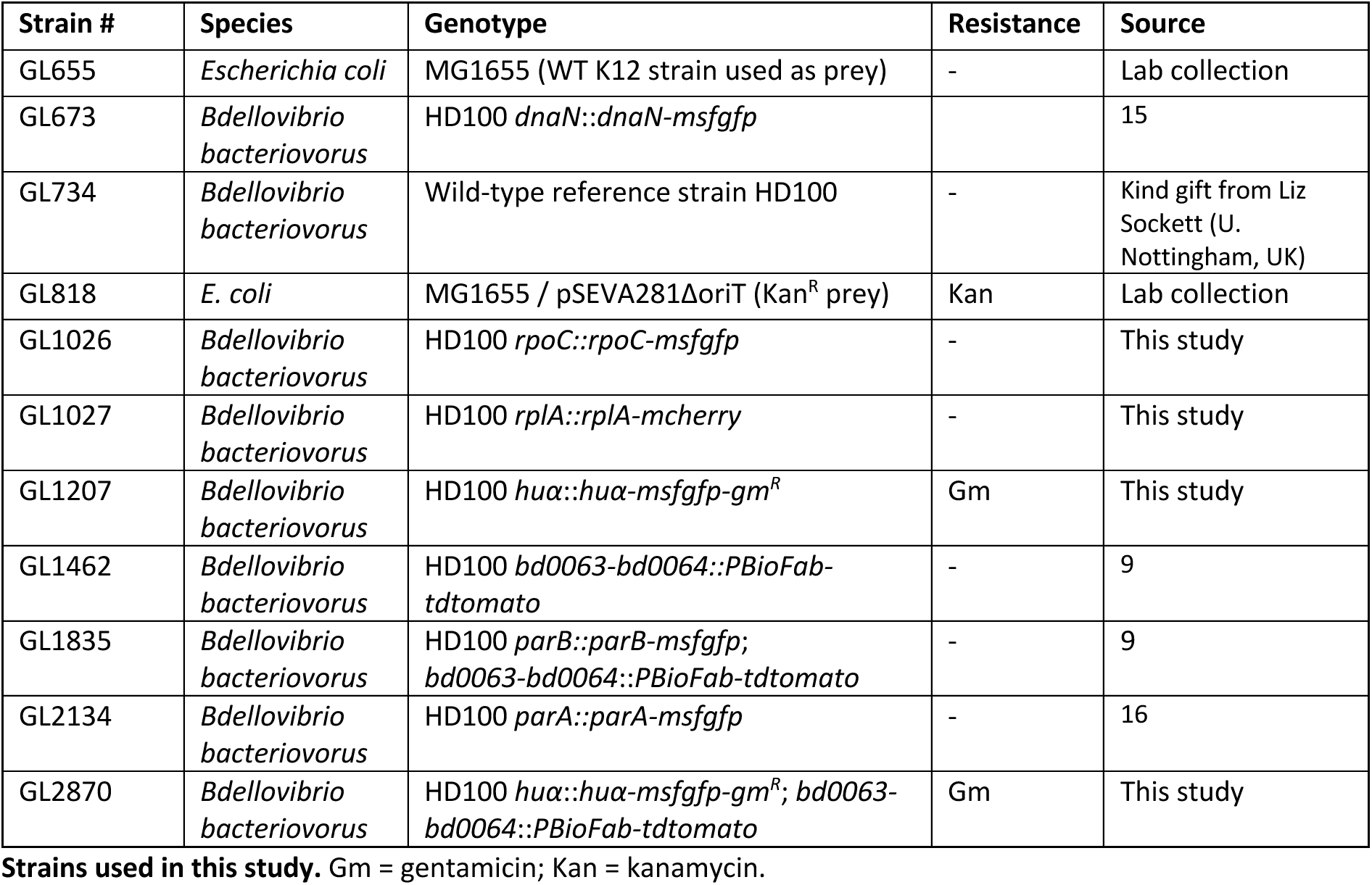

### Strains construction

Full strain construction details and oligos will be made available upon publication. Plasmids were constructed by DNA assembly using the NEB Hi-Fi Master Mix (New England Biolabs). Conjugation was performed between the *E*. *coli* S17-1 *λpir* donor strain carrying the indicated mobilizable plasmid and the *B. bacteriovorus* strain to be transformed, using the protocol described before ^15^. Plasmids were verified by sequencing of the insert. Chromosomal replacements were achieved upon a 2-step recombination strategy, and the final strain were verified by sequencing using primers hybridizing upstream and downstream the introduced modification.

### Fixation & synchronization using gradient separation

*B. bacteriovorus* attack phase cells were separated from remaining bdelloplasts and uninfected *E. coli* using an 0.8-µm filter (Millipore) and cocultured with exponentially growing *E. coli* prey to a predator:prey ratio of 1:2. 40-mL samples were taken from the *B. bacteriovorus* filtered culture and subsequently from the predation mix at 20, 50, 90, 120, 260 and 360 minutes after mixing. Immediately after sampling, formaldehyde (Merck) was added to 3% v/v. Samples were kept under mild agitation for 20 minutes at room temperature (RT) before quenching by addition of glycine (Sigma-Aldrich) to 0.5 M final concentration and incubation at RT for 15 minutes. Samples were washed three times in PBS (Sigma-Aldrich) and pelleted. The sample taken at 360 minutes was filtered on an 0.8-µm filter before fixation. The growth-phase samples (T20-T260) were resuspended in a 50 % DNB medium:Percoll (Cytiva) mixture and subsequently centrifuged for 30 minutes at max speed at RT. The top band, which appeared after separation and containing the fixed bdelloplasts, was washed 3 times in PBS and pelleted. All pellets (attack phase and growth phase) were stored at −80°C.

### RNA isolation, sequencing, and read mapping

RNA isolation was done after fixation and synchronization using the Nucleospin RNA kit (Macherey-Nagel). After RNA isolation, quality control, library preparation & sequencing was done by Novagen. Initial bioinformatic analysis was done by Novagen. In short, sequence quality control was done by Novagen using fastp and mapped to the reference genome (NCBI ASM19617v1) using Bowtie2. Quantification was done by Novagen using FeatureCounts and differential expression analysis was performed using DESeq2.

### GO enrichment analysis and gene clustering

For Gene Ontology (GO) enrichment analysis and cluster analysis, the T-Rex2 pipeline ^59^ was used using the read counts per gene generated with FeatureCounts as input. The output was further analyzed and visualized in R to create the individual time-resolved gene expression plots, the clustering heatmap and the UpSet plot.

### Preparation of samples for Hi-C

Samples were prepared for Hi-C as described in ^60^. Samples were taken out of the −80°C freezer and allowed to thaw. Each sample was then resuspended in 1.2 mL of TE buffer. Cells were lysed using the Precellys Evolution tissue homogenizer with a VK05 tube, applying five 30-second cycles, each separated by a 20-second pause. 1 mL of the lysate was transferred into a 5 mL microcentrifuge tube, supplemented with 50 µl of 10% SDS solution, and left to incubate at RT for 10 minutes. A digestion mix -consisting of 3 mL of water, 500 µl of 10% Triton, and 500 µl of 10x Digestion Buffer-was added to the solution and mixed well by pipetting. 400 µl of the sample was taken out and transferred into a 1.5 mL centrifuge tube and kept as a control (Non-Digest). 1,000 U of HpaII was added to the remaining sample and incubated at 37°C for 3 hours in a shaking incubator (∼180 rpm). Following digestion, 400 µl of the sample was taken out as a control (Digestion) and placed into a 1.5 mL microcentrifuge tube. The remaining sample was centrifuged for 20 minutes at 20°C (16,000 g). The supernatant was decanted, and the pellet was resuspended by pipetting in 400 µl of water. A biotinylation mix consisting of 50 µl 10x Ligation Buffer, 4.5 µl of 10mM dATP, dGTP, and dTTP mix, 37.5 µl of Biotin-14-dCTP, and 8 µl of DNA polymerase I (Klenow) fragment (5U/µl) was added to the resuspended pellet, mixed by pipetting, and incubated at 37°C for 45 minutes in a shaking incubator (∼180 rpm). A ligation mix consisting of 120 µl of 10x Ligation Buffer, 12 µl of 10 mg/mL BSA, 12 µl of 100 mM ATP, 540 µl of water, and 16 µl of T4 DNA ligase (30U/µl) was added to the libraries, mixed by pipetting, and incubated for 3 hours. Following the ligation reaction, a mix consisting of 20 µl 500 mM EDTA, 80 µl 10% SDS, and 100 µl 20 mg/mL proteinase K was added to each Hi-C library and mixed by pipetting. The controls, which had remained untreated, were treated with 20 µl 500 mM EDTA, 20 µl 10% SDS, and 10 µl of 20 mg/mL proteinase K. Both libraries and controls were left incubating overnight at 65°C to reverse the formaldehyde-mediated crosslinks. The next day, DNA purification by phenol extraction was performed on each Hi-C library using a 1:1 volume ratio – adding 1.2 mL of Phenol:Chloroform:Isoamyl-alcohol, vortexing for 30 seconds, removing the upper aqueous phase, and transferring it to a new 5 mL microcentrifuge tube. DNA was precipitated by adding 3 mL of 100% ethanol and 120 µl of 3M NaOAc (pH 5.0). Control DNAs were similarly precipitated by adding 1 mL of 100% ethanol and 40 µl of 3 M NaOAc (pH 5.0). Hi-C libraries and controls were left at −80°C for 30 minutes, and their DNA was pelleted by 20 minutes of centrifugation (12,000 g, 4°C). The supernatant was removed, and the pellets were dried by leaving them for 5 to 10 minutes at 37°C on a heat block. Hi-C libraries were resuspended in 140 µl of TE buffer containing 1mg/mL RNAse and left incubating for 30 min at 37°C. Controls were resuspended in 10 µl of Buffer. 10 µl of Hi-C libraries and controls were then run on an agarose gel to assess the quality of the Hi-C library. The remaining 130 µl of Hi-C libraries were transferred to a Covaris microtube and sonicated using the following parameters: Peak incidence Power 140, Duty Factor, 10%, 200 Cycles per Burst, 7° C temperature, for 80 seconds to generate a ∼300 bp size fragments. Libraries were transferred to a new microcentrifuge tube, added with an equal amount (∼130 µl) of AMPure XP beads, mixed by pipetting, and left incubating for 5 minutes to allow DNA to bind to the magnetic beads. Libraries were placed in a magnet for 5 minutes. Then, the supernatant was discarded, and the libraries were washed with 500 µl of 70% ethanol twice. Libraries were then taken off the magnet, resuspended in 300 µl of water, mixed generously, and left for 5 minutes at RT. Libraries were placed again in the magnet and the elution was transferred to a new 1.5 mL microcentrifuge tube. Libraries were mixed with 25 µl of C1 streptavidine beads resuspended in 300 µl of BB 2x buffer and allowed to incubate for 60 minutes with continuous mixing. The libraries were placed on a magnetic stand, and the supernatant was removed. The beads were washed twice with 1x TBW buffer. The libraries were then resuspended in 50 µl of TE buffer. 12.5 µl of the library was used for adding the Illumina sequencing adapters with the Colibri DNA Library Prep Kit, following the manufacturer’s instructions. The libraries were then PCR amplified using 40 µl of Phusion 2x High Fidelity MasterMix, 5 µl of 2µM Primer Mix, 3 µl of Streptavidin beads containing a Hi-C library, and 32 µl of water. The PCR ran for 12 cycles with the following parameters: initial denaturation 98°C, 30 seconds; denaturation 98°C, 10 seconds; annealing 61°C, 15 seconds; extension 72°C, 15 seconds, and final extension 72°C 10 minutes. Libraries were selected with AMPure beads as described above. Libraries concentration was quantified using the Qubit dsDNA High Sensitivity Assay Kit and prepared for paired-end sequencing.

### Hi-C analysis

Reads were checked for quality control using FastQC ^61^, then analyzed with the hicstuff pipeline ^62^.

### Microscopy

Phase contrast and fluorescence imaging was done on a Nikon Ti-2E inverted microscope (Nikon) equipped with a CFI Plan Apochromat αDM 100x 1.45/0.13 mm Ph3 oil objective (Nikon), a Sola SEII FISH illuminator (Lumencor), a temperature-controlled light-protected enclosure (Okolab) and a Photometrics Prime BSI camera. Samples for microscopy were prepared as described previously ^15^. In short, *B. bacteriovorus* and *E. coli* cells were mixed in DNB medium at time point 0 with a predator:prey ratio of 2:1 and incubated at 37°C shaking. Where applicable, cells were stained with DAPI (Life Technologies) for 5 minutes at RT before imaging. Cells were immobilized at selected time points on a 1% agarose-DNB pad before imaging. Illumination and exposure times were kept identical when imaging different strains and/or conditions in one experiment.

### Image processing

Images were processed with FIJI^63^ keeping the brightness and contrast settings identical for all regions of interest in each figure. Figures were assembled and annotated using Adobe Indesign (Adobe).

### Segmentation & measurements using MiSiC

For measuring the cell and nucleoid size during a full time-course, fluorescent images were denoised using the Noise2Void algorithm ^41^. A MiSiC model ^40^ was trained on fluorescence images of growing *B. bacteriovorus* producing cytoplasmic tdTomato and manually curated for multiple learning rounds. *B. bacteriovorus* cells were segmented using images of *B. bacteriovorus* producing cytoplasmic tdTomato as input for MiSiC, nucleoids were segmented using images of *B. bacteriovorus* producing HUα-sfGFP and/or DAPI-stained nucleoids as input for MiSiC. The resulting binary masks were checked by eye and *B. bacteriovorus* cells that were not inside a bdelloplast were removed from analysis. The binary masks were used as input for MicrobeJ ^64^ to obtain straightened fluorescence profiles and nucleoid/cell length, area and width. Using the R package BactMAP ^65^, MicrobeJ contour output and TIFF images were transformed to dataframes and cell data was related to their nucleoid data. Using this, the nucleocytoplasmic ratio was calculated. The straightened fluorescence profiles were used to calculate the median intensity profiles (**Figure 2**). The median intensity over the length axis was calculated from the fluorescence profiles, and the difference between the DAPI and tdTomato profiles was calculated by normalizing both profiles and subsequently subtracting the DAPI from the tdTomato signal.

### Nucleocytoplasmic ratio measurement in attack phase cells

For **Figure 3EF**, only AP cells were measured. Here, cells and nucleoids were segmented as described previously ^15^. In short, Oufti *celldetection* tool ^66^ was used to segment the cells from phase contrast images and *objectdetection* was used to segment the DAPI signal. Subsequently, the object and cell information were transformed to dataframes using BactMAP ^65^, after which ggplot2 ^67^ was used to plot the nucleocytoplasmic ratio of each sample. TIFFs were transformed to dataframes using BactMAP to subsequently plot the median fluorescence profiles and cell shapes.

### MIC determination

The number of *B. bacteriovorus* cells was determined upon SyberGreen staining as described before ^33^. *Bdellovibrio bacteriovorus* and *E. coli* MG1655 were mixed to a predator:prey ration of 1:1 in DNB-salts with a titration of chloramphenicol or rifampicin concentrations (see **Figure S8**) in technical triplicates. Killing curves were measured as described previously in a Synergy H1m microplate reader (Biotek) ^9,33^. OD_600_ and fluorescence at 568 nm were measured every 20 minutes under continuous shaking at 30°C. The experiment was carried out in duplicate. The P_max_ of each curve was determined using the R package CuRveR ^33^, and the MIC curves were plotted in R using the package ggplot2.

### Sample preparation for ChIP-sequencing

Samples of GL734 (WT, mock control), GL1026 (RpoC-msfGFP) and GL2134 (ParA-msfGFP) were taken from overnight cultures (T0) and 2 hours after co-incubation with *E. coli* MG1655 cells (T120) in DNB-salts. Overnight cultures were prepared by inoculating 1 mL of *B. bacteriovorus* lysate and 5 mL of *E. coli* prey (in Ca-HEPES, OD₆₀₀ ≈ 10) into 44 mL DNB-salts, yielding 50 mL per replicate. Cultures were incubated overnight at 30°C with shaking. The following day, AP cells were isolated by filtration through a 1.2-µm filter (Millipore). From each filtered culture, 40 mL was used for fixation, and 10 mL was used to initiate growth-phase (GP) cultures. To fix AP cells, formaldehyde (Merck) was added to the AP samples to a final concentration of 1% (v/v), followed by incubation at RT with gentle agitation for 30 minutes. Crosslinking was quenched with 2.5 mL of 2.5 M glycine (Sigma-Aldrich) for 15 minutes at RT. Cells were pelleted and washed three times in 1× PBS (10 minutes, 6000 rpm, RT). GP cultures were initiated by combining 10 mL of filtered AP lysate, 5 mL of *E. coli* prey (OD_600_ ≈ 10), and 35 mL DNB-salts per replicate. Cultures were incubated at 30°C for 2 hours. After 2h of predation, fixation was performed as described above, without prior filtration. Fixed GP cells were resuspended in a 1:1 mixture of DNB medium and Percoll (Cytiva), then centrifuged at maximum speed for 30 minutes at RT. The upper yellow band containing fixed bdelloplasts was carefully aspirated and transferred to fresh tubes. Cells were washed three times in 1× PBS and pelleted (10 minutes, 6000 rpm, RT). All AP and GP pellets were stored at −80°C until further processing.

### ChIP and library preparation

Cell pellets were resuspended in 1 mL buffer 1 (20 mM K-HEPES pH 7.9, 50 mM KCl, 10% glycerol, and EDTA-free protease inhibitors (Roche)). Chromatin was sheared by sonication on ice using a Soniprep 150 (15 cycles: 15 s ON, 15 s OFF, setting 10), and debris was removed by centrifugation (20 minutes, 17,000 × g, 4°C). The supernatant was transferred to fresh tubes, and buffer conditions were adjusted to 10 mM Tris-HCl, pH 8, 150 mM NaCl, and 0.1% NP-40. In parallel, 20 μL GFP antibody-conjugated beads (Abcam) were washed with IPP150 buffer (10 mM Tris-HCl pH 8, 150 mM NaCl, 0.1% NP-40) and added to the lysates. Samples were incubated overnight at 4°C with gentle rotation. The following day, beads were washed five times with 1 mL IPP150 buffer and twice with 1× TE buffer (10 mM Tris-HCl pH 8, 1 mM EDTA) at 4°C. Protein–DNA complexes were eluted in two steps: first with 150 μL elution buffer (50 mM Tris-HCl pH 8, 10 mM EDTA, 1% SDS) for 15 minutes at 65°C, then with 100 μL 1× TE buffer containing 1% SDS under the same conditions. The supernatant (i.e., the ChIP fraction) was then separated from the beads and further incubated at 65 °C overnight to reverse crosslinks completely. DNA was purified using the QIAquick PCR Purification Kit (Qiagen) and eluted in 40 μL nuclease-free water. ChIP-seq libraries were prepared using the NEBNext Ultra II DNA Library Prep Kit (NEB) and sequenced on the Illumina Hiseq 2500 or Nextseq 550 at the Tufts University Genomics facility.

### ChIP-sequencing analysis

For the analysis, sequence trimming, alignment, and peak annotation were done using a snakemake pipeline ^68^. Sequence files were gunzipped, and sequence quality was assessed using FastQC^61^. Data was trimmed using Trimmomatic ^69^, and sequences were aligned to the reference genome (NCBI ASM19617v1) using Bowtie 2.0 ^70^ and converted to .bam files using SamTools ^71^ and filtered using Sambamba ^72^. Sequence read depths were binned by 50 bp, normalized, averaged, and IP over input was calculated using Deeptools.

## DATA AVAILABILITY

Codes, sequencing data, and supplementary datasets are currently under controlled access and will be openly released upon publication.

## AUTHOR CONTRIBUTIONS

RR and GL designed experiments and wrote the first drafts. RR performed experiments and analyzed the data. FOYC performed Hi-C, mapped, and visualized the Hi-C data, with contributions from CP. JK performed sampling for the ChIP-sequencing. NT performed ChIP-sequencing under supervision of TBKL. SP trained MiSiC for *B. bacteriovorus.* RK supervised the Hi-C analyses. GL wrote the final version of the manuscript, supervised the work, and obtained funding.

## ACKNOWLEDGEMENTS

We are grateful to all members of the Laloux lab for insightful discussions and thorough reading of the manuscript; Coralie Tesseur for the initial schematics of the *B. bacteriovorus* lifecycle; and former lab members Terrens Saaki and Antoine Gérard for constructing strain GL1207, and strains GL1026 and GL1027, respectively.

## REFERENCES

1. Reyes-Lamothe, R. & Sherratt, D. J. The bacterial cell cycle, chromosome inheritance and cell growth. Nat Rev Micro 15, 1 (2019).

2. Wang, X., Llopis, P. M. & Rudner, D. Z. Organization and segregation of bacterial chromosomes. Nat. Rev. Genet. 14, 191–203 (2013).

3. Abbondanzieri, E. A. et al. Future Directions of the Prokaryotic Chromosome Field. Mol. Microbiol. 123, 89–100 (2025).

4. Tesseur, C., Santin, Y. G. & Laloux, G. Strategies and mechanisms of contact-dependent predation in bacteria. Curr. Opin. Microbiol. 87, 102639 (2025).

5. Laloux, G. Shedding Light on the Cell Biology of the Predatory Bacterium Bdellovibrio bacteriovorus. Front. Microbiol. 10, 3136 (2020).

6. Lai, T. F., Ford, R. M. & Huwiler, S. G. Advances in cellular and molecular predatory biology of Bdellovibrio bacteriovorus six decades after discovery. Front. Microbiol. 14, 1168709 (2023).

7. Caulton, S. G. & Lovering, A. L. Moving toward a better understanding of the model bacterial predator Bdellovibrio bacteriovorus. Microbiology 169, 001380 (2023).

8. Kuru, E. et al. Fluorescent D-amino-acids reveal bi-cellular cell wall modifications important for Bdellovibrio bacteriovorus predation. Nature Microbiology 2, 1648–1657 (2017).

9. Santin, Y. G., Lamot, T., Raaphorst, R. van, Kaljević, J. & Laloux, G. Modulation of prey size reveals adaptability and robustness in the cell cycle of an intracellular predator. Curr. Biol. 33, 2213–2222.e4 (2023).

10. Kessel, M. & Shilo, M. Relationship of Bdellovibrio elongation and fission to host cell size. J Bacteriol 128, 477–480 (1976).

11. Deghelt, M. et al. G1-arrested newborn cells are the predominant infectious form of the pathogen Brucella abortus. Nat. Commun. 5, 4366 (2014).

12. Salje, J. Cells within cells: Rickettsiales and the obligate intracellular bacterial lifestyle. Nat Rev Micro 151, 4015 (2021).

13. Figueroa-Cuilan, W. M. et al. Quantitative analysis of morphogenesis and growth dynamics in an obligate intracellular bacterium. Mol. Biol. Cell 34, ar69 (2023).

14. Karunker, I., Rotem, O., Dori-Bachash, M., Jurkevitch, E. & Sorek, R. A Global Transcriptional Switch between the Attack and Growth Forms of Bdellovibrio bacteriovorus. Plos One 8, e61850 (2013).

15. Kaljević, J. et al. Chromosome choreography during the non-binary cell cycle of a predatory bacterium. Curr. Biol. 31, 3707–3720.e5 (2021).

16. Kaljević, J., Tesseur, C., Le, T. B. K. & Laloux, G. Cell cycle-dependent organization of a bacterial centromere through multi-layered regulation of the ParABS system. PLOS Genet. 19, e1010951 (2023).

17. Makowski, Ł. et al. Dynamics of chromosome replication and its relationship to predatory attack lifestyles in Bdellovibrio bacteriovorus. Appl. Environ. Microbiol. (2019) doi:10.1128/aem.00730-19.

18. Hocher, A. et al. Histones with an unconventional DNA-binding mode in vitro are major chromatin constituents in the bacterium Bdellovibrio bacteriovorus. Nat. Microbiol. 1–14 (2023) doi:10.1038/s41564-023-01492-x.

19. Hu, Y. et al. Bacterial histone HBb from Bdellovibrio bacteriovorus compacts DNA by bending. Nucleic Acids Res. gkae485 (2024) doi:10.1093/nar/gkae485.

20. Yáñez-Cuna, F. O. & Koszul, R. Insights in bacterial genome folding. Curr. Opin. Struct. Biol. 82, 102679 (2023).

21. Gaal, T. et al. Colocalization of distant chromosomal loci in space in E. coli: a bacterial nucleolus. Genes Dev. 30, 2272–2285 (2016).

22. Ladouceur, A.-M. et al. Clusters of bacterial RNA polymerase are biomolecular condensates that assemble through liquid–liquid phase separation. Proc. Natl. Acad. Sci. 117, 18540–18549 (2020).

23. Stracy, M. et al. Live-cell superresolution microscopy reveals the organization of RNA polymerase in the bacterial nucleoid. Proc. Natl. Acad. Sci. USA 112, E4390–9 (2015).

24. Chai, Q. et al. Organization of Ribosomes and Nucleoids in Escherichia coli Cells during Growth and in Quiescence. J. Biol. Chem. 289, 11342–11352 (2014).

25. Dworsky, P. & Schaechter, M. Effect of Rifampin on the Structure and Membrane Attachment of the Nucleoid of Escherichia coli. J. Bacteriol. 116, 1364–1374 (1973).

26. Bakshi, S., Choi, H., Mondal, J. & Weisshaar, J. C. Time-dependent effects of transcription- and translation-halting drugs on the spatial distributions of the Escherichia coli chromosome and ribosomes. Mol. Microbiol. 94, 871–887 (2014).

27. Bignaud, A. et al. Transcription-induced domains form the elementary constraining building blocks of bacterial chromosomes. Nat. Struct. Mol. Biol. 31, 489–497 (2024).

28. Le, T. B. & Laub, M. T. Transcription rate and transcript length drive formation of chromosomal interaction domain boundaries. EMBO J. 35, 1582–1595 (2016).

29. Amemiya, H. M., Schroeder, J. & Freddolino, P. L. Nucleoid-associated proteins shape chromatin structure and transcriptional regulation across the bacterial kingdom. Transcription 12, 182–218 (2021).

30. Melfi, M. D., Shapiro, L., Lasker, K. & Zhou, X. ATAC-seq reveals megabase-scale domains of a bacterial nucleoid. bioRxiv 12, 2021.01.09.426053 (2021).

31. Lioy, V. S. et al. Multiscale Structuring of the E. coli Chromosome by Nucleoid-Associated and Condensin Proteins. Cell 172, 771–783.e18 (2018).

32. Lambert, C., Chang, C.-Y., Capeness, M. J. & Sockett, R. E. The first bite--profiling the predatosome in the bacterial pathogen Bdellovibrio. PLoS ONE 5, e8599 (2010).

33. Remy, O. et al. An optimized workflow to measure bacterial predation in microplates. STAR Protoc. 3, 101104 (2022).

34. Rosson, R. A. & Rittenberg, S. C. Regulated breakdown of Escherichia coli deoxyribonucleic acid during intraperiplasmic growth of Bdellovibrio bacteriovorus 109J. J. Bacteriol. 140, 620–33 (1979).

35. Yang, T. et al. HiCRep: assessing the reproducibility of Hi-C data using a stratum-adjusted correlation coefficient. Genome Res. 27, 1939–1949 (2017).

36. Marbouty, M. et al. Condensin- and Replication-Mediated Bacterial Chromosome Folding and Origin Condensation Revealed by Hi-C and Super-resolution Imaging. Mol. Cell 59, 588–602 (2015).

37. Wang, X., Brandão, H. B., Le, T. B. K., Laub, M. T. & Rudner, D. Z. Bacillus subtilis SMC complexes juxtapose chromosome arms as they travel from origin to terminus. Science 355, 524–527 (2016).

38. Tran, N. T., Laub, M. T. & Le, T. B. K. SMC progressively aligns chromosomal arms in caulobacter crescentus but is antagonized by convergent transcription. Cell Reports 20, 2057–2071 (2017).

39. Badrinarayanan, A., Reyes-Lamothe, R., Uphoff, S., Leake, M. C. & Sherratt, D. J. In Vivo Architecture and Action of Bacterial Structural Maintenance of Chromosome Proteins. Science 338, 528–531 (2012).

40. Panigrahi, S. et al. Misic, a general deep learning-based method for the high-throughput cell segmentation of complex bacterial communities. Elife 10, e65151 (2021).

41. Krull, A., Buchholz, T.-O. & Jug, F. Noise2Void - Learning Denoising from Single Noisy Images. 2019 IEEECVF Conf. Comput. Vis. Pattern Recognit. (CVPR) 00, 2124–2132 (2019).

42. Bettridge, K., Verma, S., Weng, X., Adhya, S. & Xiao, J. Single-molecule tracking reveals that the nucleoid-associated protein HU plays a dual role in maintaining proper nucleoid volume through differential interactions with chromosomal DNA. Mol. Microbiol. 115, 12–27 (2021).

43. Cabrera, J. E., Cagliero, C., Quan, S., Squires, C. L. & Jin, D. J. Active transcription of rRNA operons condenses the nucleoid in Escherichia coli: examining the effect of transcription on nucleoid structure in the absence of transertion. J. Bacteriol. 191, 4180–4185 (2009).

44. Lambert, C. et al. Characterizing the flagellar filament and the role of motility in bacterial prey-penetration by Bdellovibrio bacteriovorus. Mol. Microbiol. 60, 274–286 (2006).

45. Remy, O. et al. Distinct dynamics and proximity networks of hub proteins at the prey-invading cell pole in a predatory bacterium. J. Bacteriol. 206, e00014–24 (2024).

46. Kaplan, M. et al. Bdellovibrio predation cycle characterized at nanometre-scale resolution with cryo-electron tomography. Nat. Microbiol. 1–13 (2023) doi:10.1038/s41564-023-01401-2.

47. Lerner, T. R. et al. Specialized peptidoglycan hydrolases sculpt the intra-bacterial niche of predatory Bdellovibrio and increase population fitness. PLoS Pathog. 8, e1002524 (2012).

48. Banks, E. J. et al. An MltA-Like Lytic Transglycosylase Secreted by Bdellovibrio bacteriovorus Cleaves the Prey Septum during Predatory Invasion. J Bacteriol e00475–22 (2023) doi:10.1128/jb.00475-22.

49. Harding, C. J. et al. A lysozyme with altered substrate specificity facilitates prey cell exit by the periplasmic predator Bdellovibrio bacteriovorus. Nat Comms 11, 4817 (2020).

50. Lambert, C. et al. Interrupting peptidoglycan deacetylation during Bdellovibrio predator-prey interaction prevents ultimate destruction of prey wall, liberating bacterial-ghosts. Sci. Rep. 6, 26010 (2016).

51. Banks, E. J. et al. Asymmetric peptidoglycan editing generates cell curvature in Bdellovibrio predatory bacteria. Nat Commun 13, 1509 (2022).

52. Lai, T. F., Jankov, D., Grossmann, J., Roschitzki, B. & Huwiler, S. G. Quantitative proteome of bacterial periplasmic predation reveals a prey damaging protease. bioRxiv 2024.12.23.630089 (2024) doi:10.1101/2024.12.23.630089.

53. Le, T. B. K., Imakaev, M. V., Mirny, L. A. & Laub, M. T. High-Resolution Mapping of the Spatial Organization of a Bacterial Chromosome. Science 342, 731–734 (2013).

54. Szafran, M. J. et al. Spatial rearrangement of the Streptomyces venezuelae linear chromosome during sporogenic development. Nat Comms 12, 1–15 (2021).

55. Lioy, V. S. et al. Dynamics of the compartmentalized Streptomyces chromosome during metabolic differentiation. Nat Comms 12, 1–14 (2021).

56. Spahn, C. et al. The nucleoid of rapidly growing Escherichia coli localizes close to the inner membrane and is organized by transcription, translation, and cell geometry. Nat. Commun. 16, 3732 (2025).

57. Gadkari, D. & Stolp, H. Energy metabolism of Bdellovibrio bacteriovorus. Arch. Microbiol. 102, 179–185 (1975).

58. Herencias, C., Salgado-Briegas, S., Prieto, M. A. & Nogales, J. Providing new insights on the biphasic lifestyle of the predatory bacterium Bdellovibrio bacteriovorus through genome-scale metabolic modeling. PLoS Comput. Biol. 16, e1007646 (2020).

59. Jong, A. de, Meulen, S. van der, Kuipers, O. P. & Kok, J. T-REx: Transcriptome analysis webserver for RNA-seq Expression data. BMC Genom. 16, 663 (2015).

60. Cockram, C., Thierry, A. & Koszul, R. Generation of gene-level resolution chromosome contact maps in bacteria and archaea. STAR Protocols 2, 100512 (2021).

61. Andrews, S. FastQC: A Quality Control Tool for High Throughput Sequence Data. (2010).

62. Matthey-Doret, C. et al. Hicstuff: Simple Library/Pipeline to Generate and Handle Hi-C Data. (2020).

63. Schindelin, J., et al. Fiji: an open-source platform for biological-image analysis. Nat. Methods 9, 676–682 (2012).

64. Ducret, A., Quardokus, E. M. & Brun, Y. V. MicrobeJ, a tool for high throughput bacterial cell detection and quantitative analysis. Nature Microbiology 1, 16077 (2016).

65. Raaphorst, R. van, Kjos, M. & Veening, J.-W. BactMAP: An R package for integrating, analyzing and visualizing bacterial microscopy data. Mol. Microbiol. 113, 297–308 (2020).

66. Paintdakhi, A. et al. Oufti: an integrated software package for high-accuracy, high-throughput quantitative microscopy analysis. Mol. Microbiol. 99, 767–777 (2016).

67. Wickham, H. Ggplot2: Elegant Graphics for Data Analysis. (Springer-Verlag New York, 2016). doi:10.1007/978-0-387-98141-3.

68. Mazzuoli, M.-V. et al. HU promotes higher-order chromosome organisation and influences DNA replication rates in Streptococcus pneumoniae. bioRxiv 2024.09.27.615122 (2024) doi:10.1101/2024.09.27.615122.

69. Bolger, A. M., Lohse, M. & Usadel, B. Trimmomatic: a flexible trimmer for Illumina sequence data. Bioinformatics 30, 2114–2120 (2014).

70. Langmead, B. & Salzberg, S. L. Fast gapped-read alignment with Bowtie 2. Nat. Methods 9, 357–359 (2012).

71. Li, H. et al. The Sequence Alignment/Map format and SAMtools. Bioinformatics 25, 2078–2079 (2009).

72. Tarasov, A., Vilella, A. J., Cuppen, E., Nijman, I. J. & Prins, P. Sambamba: fast processing of NGS alignment formats. Bioinformatics 31, 2032–2034 (2015).

