## Supplementary Figures and Legends for "Dynamics and interplay of gene expression and chromosome organization across a predatory lifecycle"

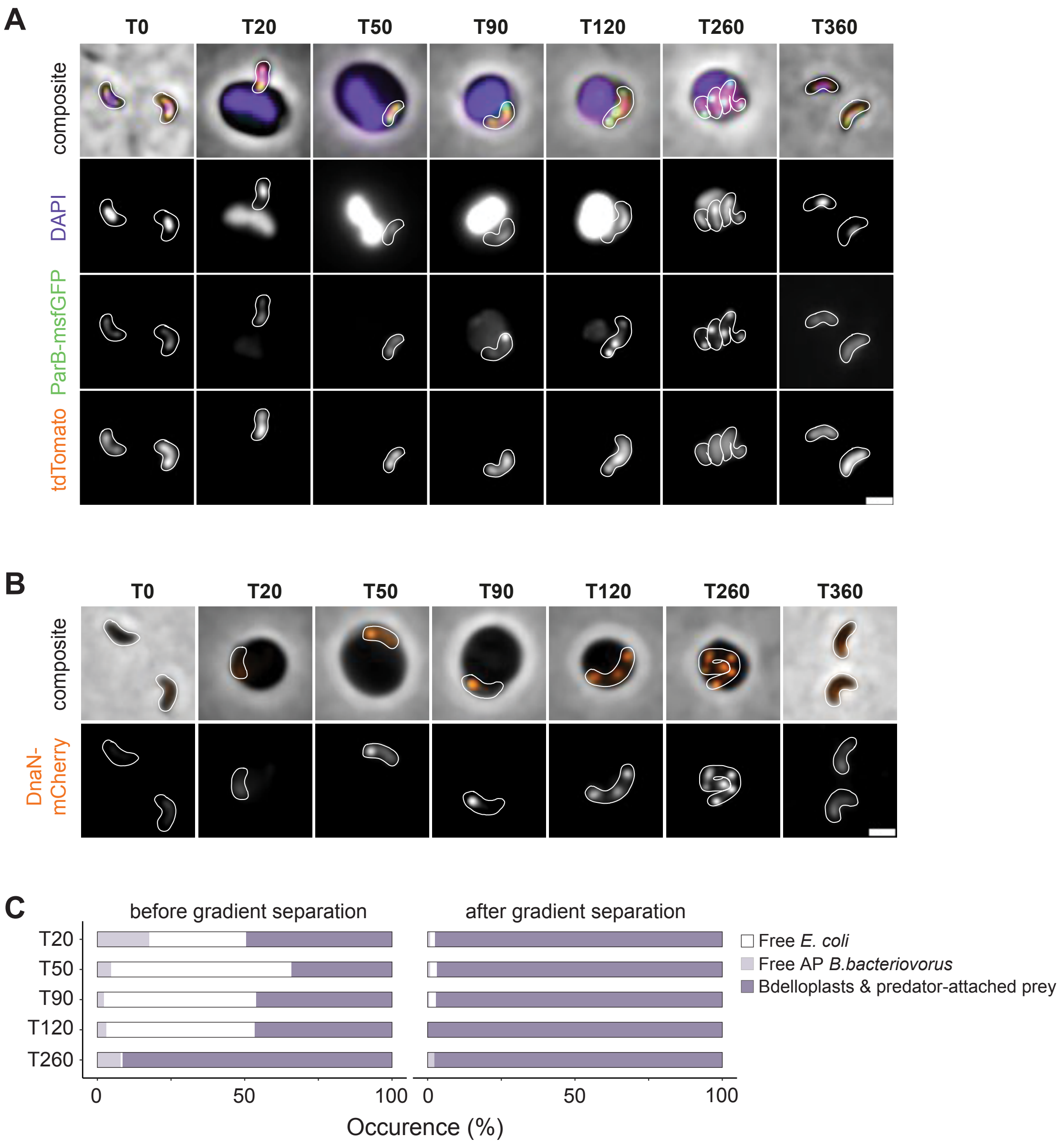

A
T0
T20
T50
T90
T120
T260
T360
composite
DAPI
ParB-msfGFP
tdTomato
B
DnaN-mCherry
C
before gradient separation
after gradient separation
T20
T50
T90
T120
T260
0
50
100
Occurence (%)
Free E. coli
Free AP B.bacteriovorus
Bdelloplasts & predator-attached prey


Figure S1

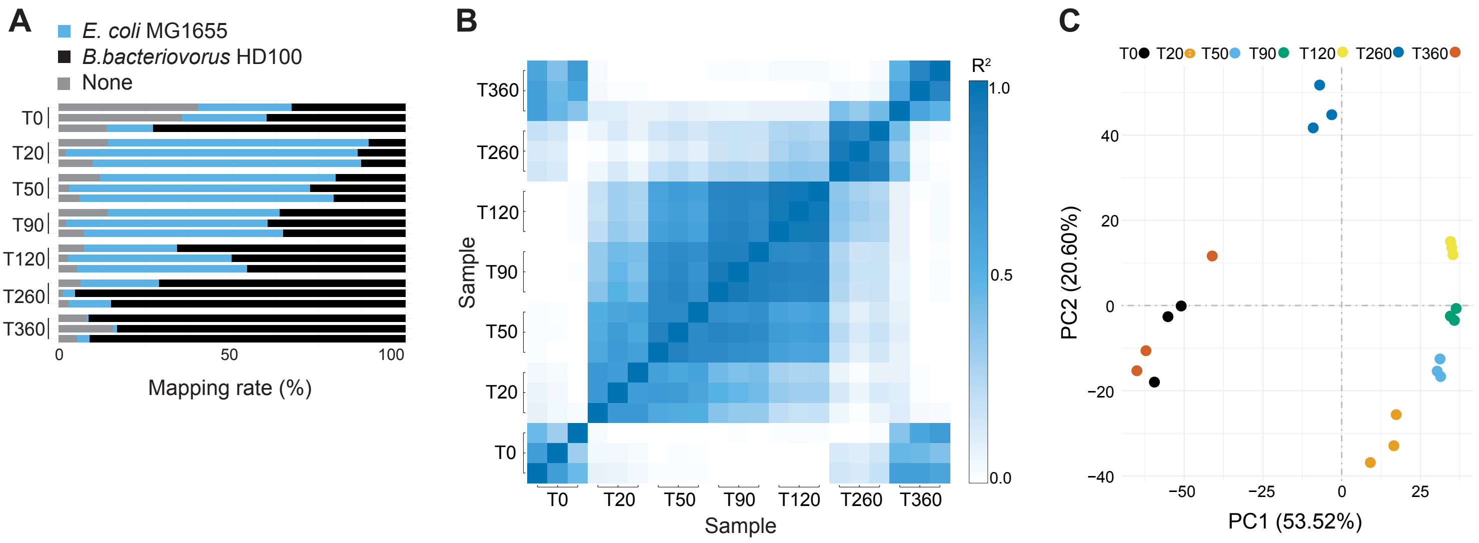
A
E. coli MG1655
B.bacteriovorus HD100
None
T0
T20
T50
T90
T120
T260
T360
0
50
100
Mapping rate (%)
B
Sample
R²
1.0
0.5
0.0
C
T0 T20 T50 T90 T120 T260 T360
PC2 (20.60%)
PC1 (53.52%)
40
20
0
−20
−40
−50
−25
25

Figure S2

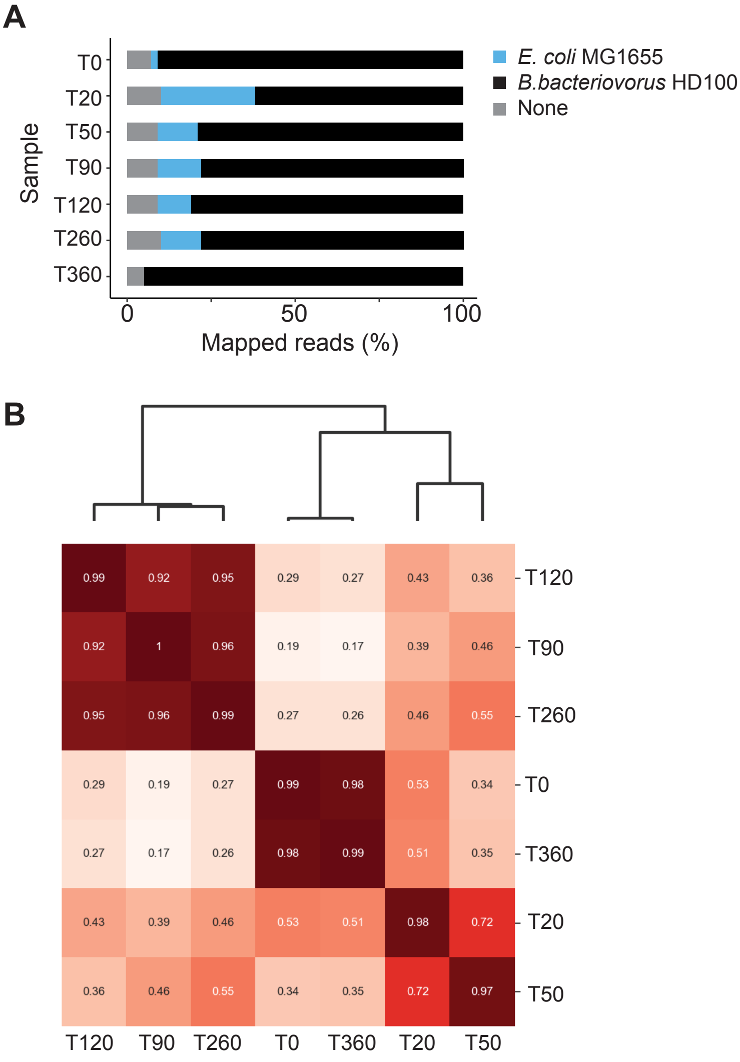
A
Sample
T0
T20
T50
T90
T120
T260
T360
0
50
100
Mapped reads (%)
E. coli MG1655
B.bacteriovorus HD100
None
B
T120
T90
T260
T0
T360
T20
T50
0.99 0.92 0.95 0.29 0.27 0.43 0.36
0.92 1 0.96 0.19 0.17 0.39 0.46
0.95 0.96 0.99 0.27 0.26 0.46 0.55
0.29 0.19 0.27 0.99 0.98 0.53 0.34
0.27 0.17 0.26 0.98 0.99 0.51 0.35
0.43 0.39 0.46 0.53 0.51 0.98 0.72
0.36 0.46 0.55 0.34 0.35 0.72 0.97
T120 T90 T260 T0 T360 T20 T50

Figure S3

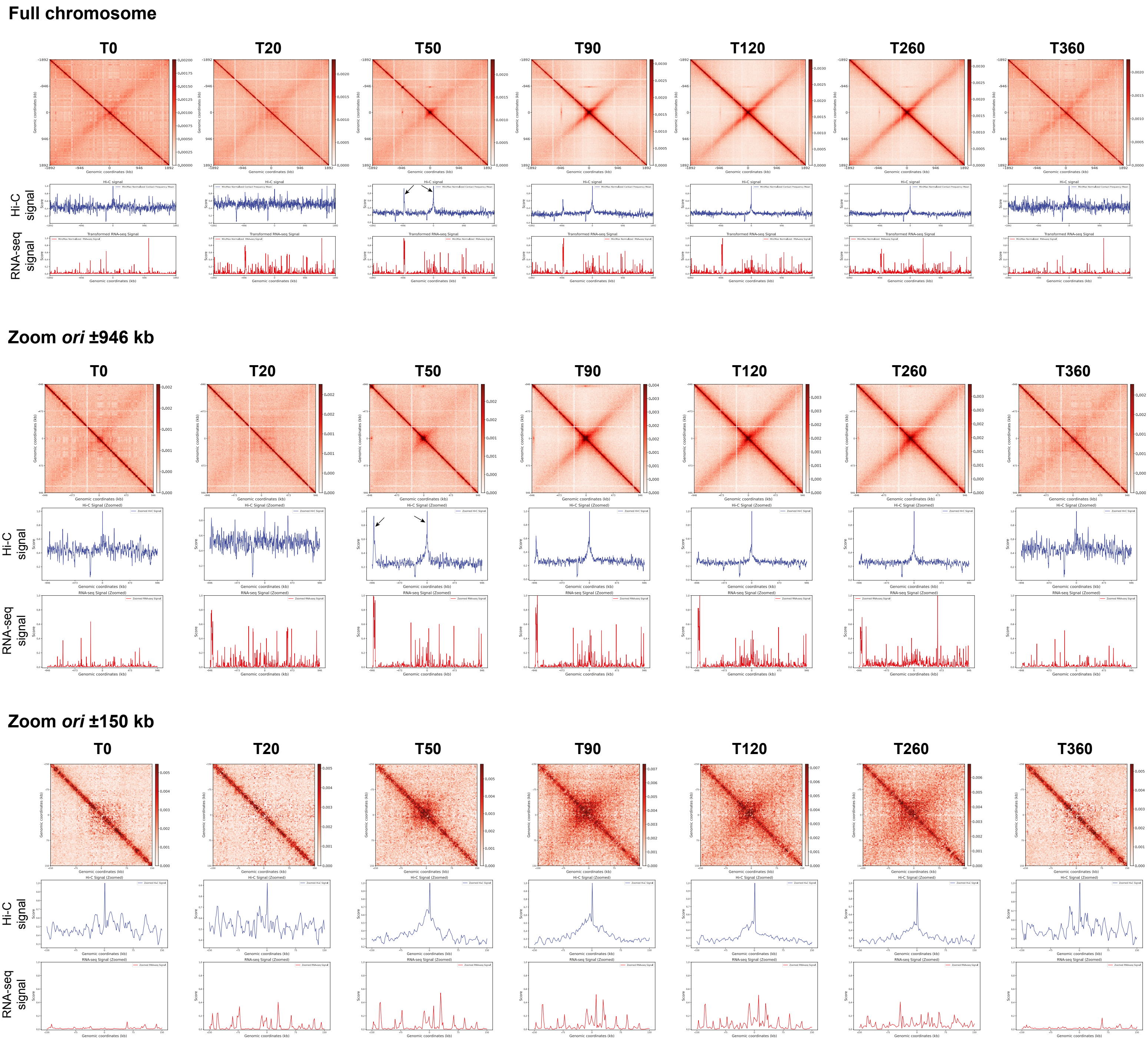
Full chromosome
T0
T20
T50
T90
T120
T260
T360
Hi-C signal
RNA-seq signal
Zoom ori ±946 kb
Zoom ori ±150 kb
Genomic coordinates (kb)
Hi-C signal
Transformed RNA-seq Signal
Hi-C Signal (Zoomed)
RNA-seq Signal (Zoomed)

Figure S4

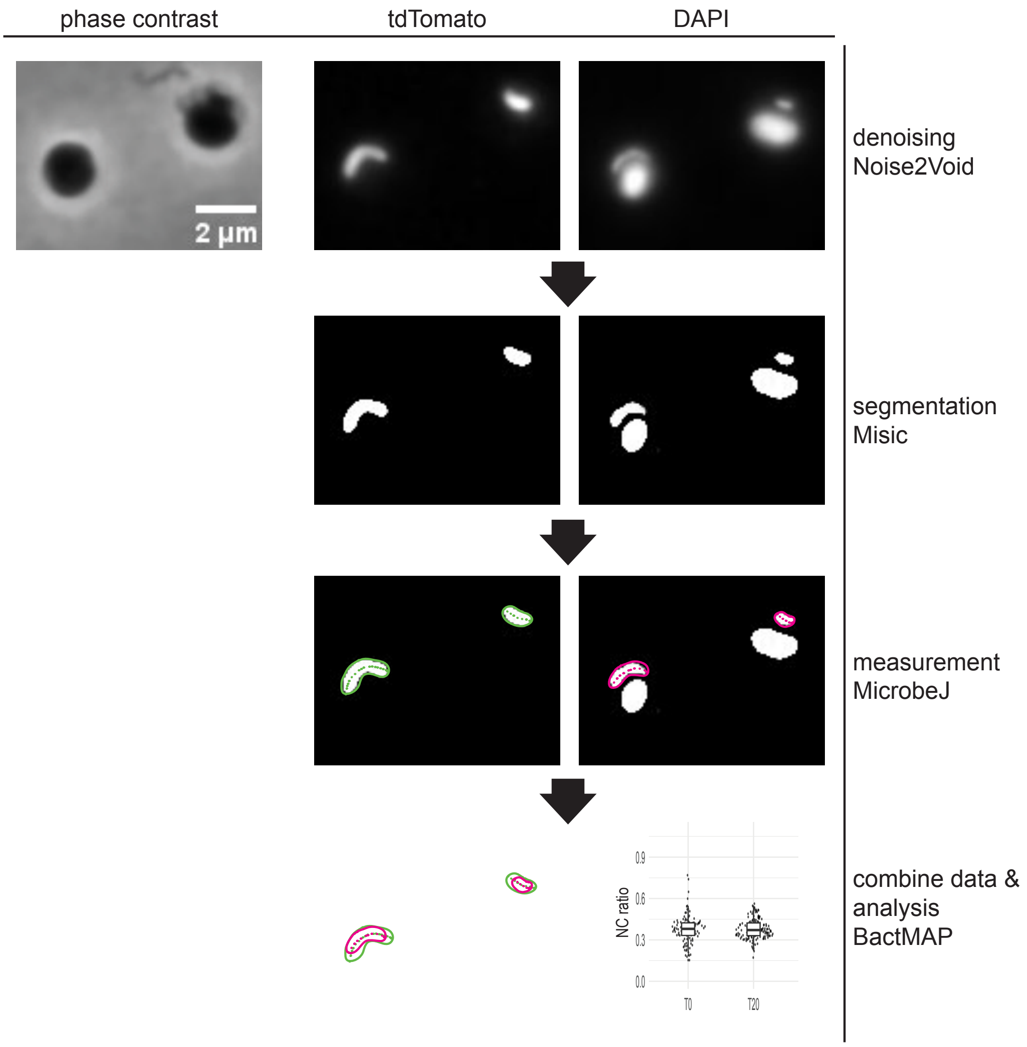
phase contrast
tdTomato
DAPI
2 µm
denoising
Noise2Void
segmentation
Misic
measurement
MicrobeJ
combine data &
analysis
BactMAP
NC ratio
0.9
0.6
0.3
0.0
T0
T20

Figure S5

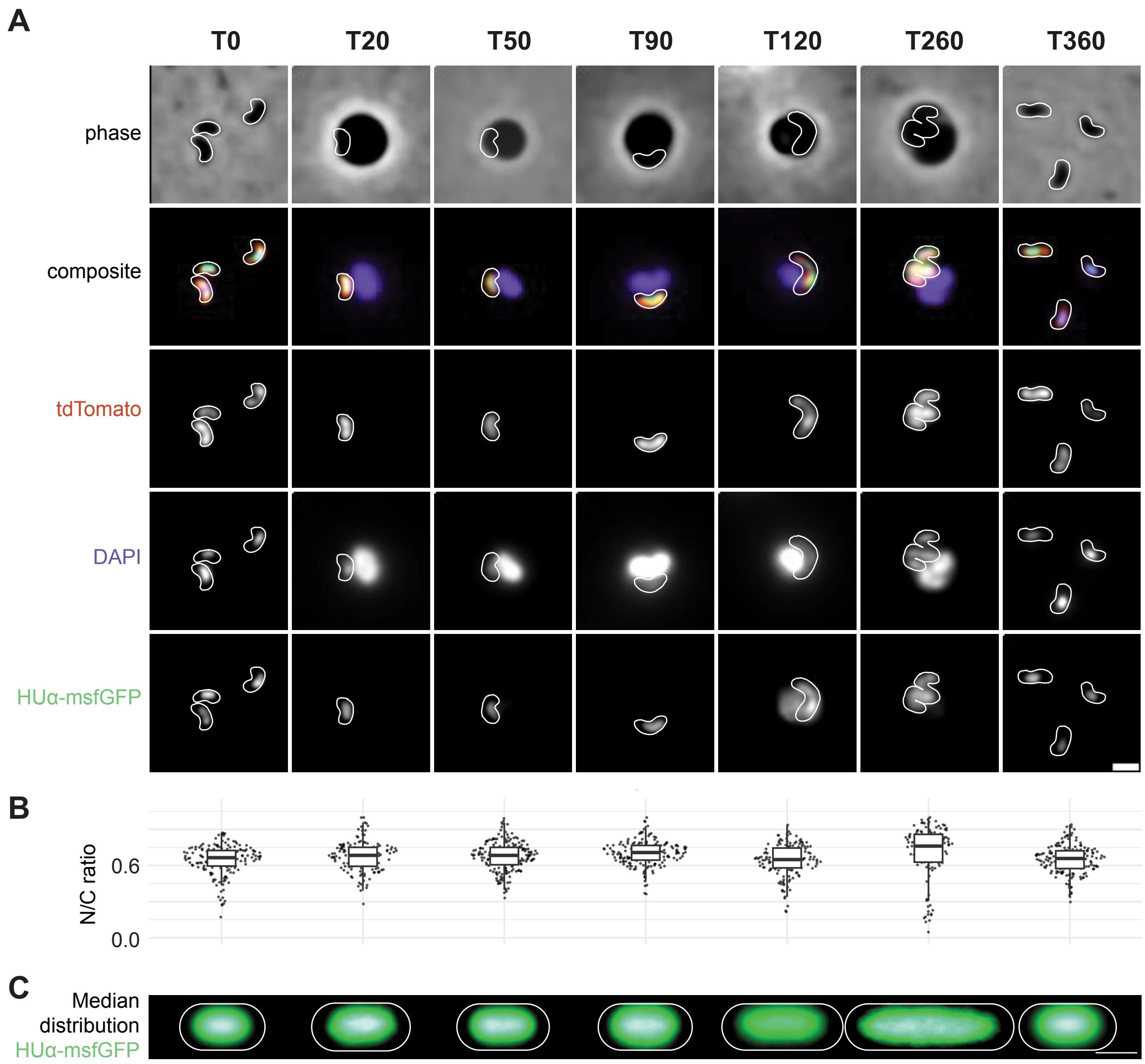
A
T0
T20
T50
T90
T120
T260
T360
phase
composite
tdTomato
DAPI
HUα-msfGFP
B
N/C ratio
0.6
0.0
C
Median distribution HUα-msfGFP

Figure S6

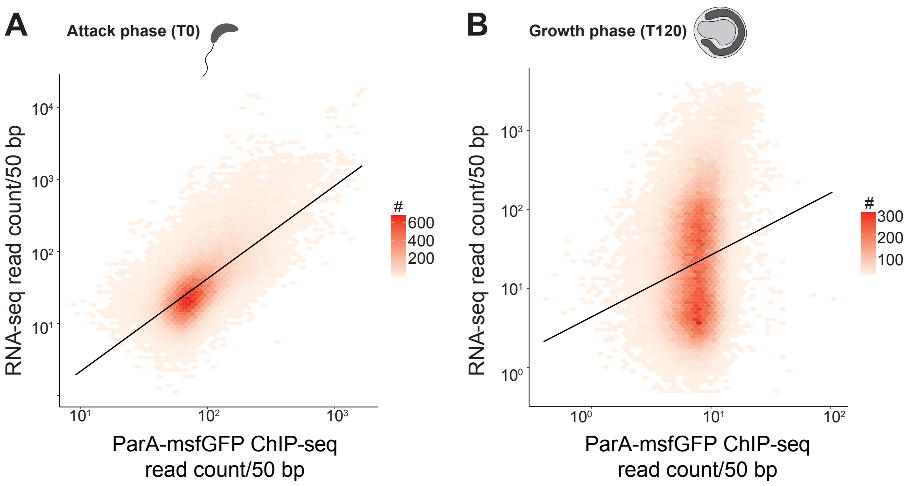
A
Attack phase (T0)
RNA-seq read count/50 bp
ParA-msfGFP ChIP-seq read count/50 bp
#
600
400
200
B
Growth phase (T120)
RNA-seq read count/50 bp
ParA-msfGFP ChIP-seq read count/50 bp
#
300
200
100

Figure S7

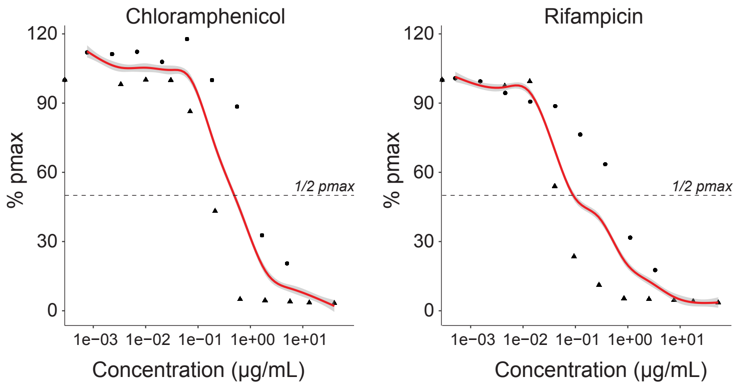

Chloramphenicol
Rifampicin
% pmax
120
90
60
30
0
1/2 pmax
1e−03
1e−02
1e−01
1e+00
1e+01
Concentration (μg/mL)


Figure S8

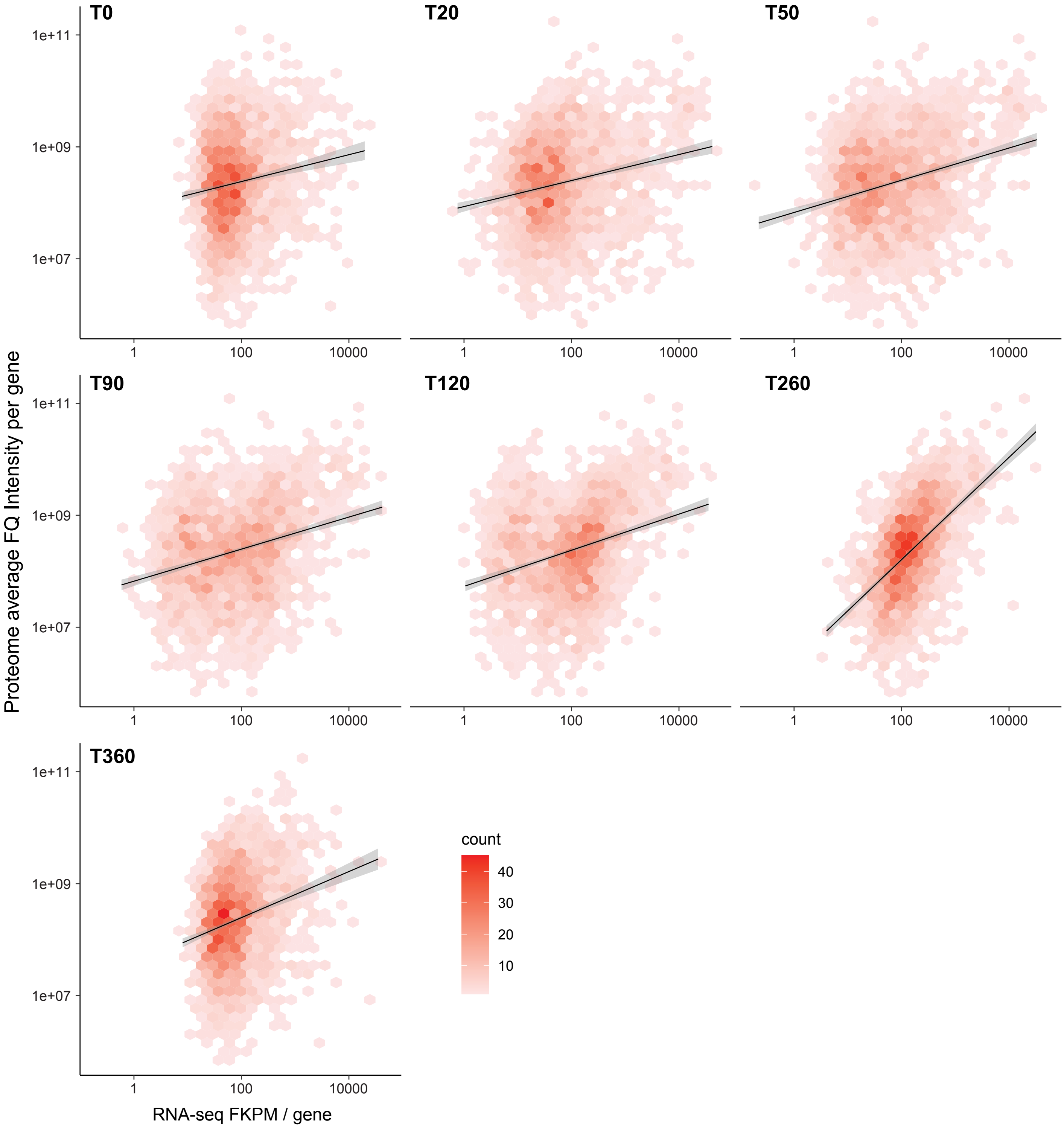
T0
T20
T50
T90
T120
T260
T360
1e+11
1e+09
1e+07
1
100
10000
Proteome average FQ Intensity per gene
RNA-seq FKPM / gene
count
40
30
20
10

Figure S9

### SUPPLEMENTARY FIGURE LEGENDS

**Figure S1. A.** Representative images of *B. bacteriovorus* (strain GL1835) expressing *parB-msfgfp* from the native *parB* locus and constitutively producing cytoplasmic tdTomato, showing the first appearance of a ParB focus at 90 minutes after co-incubating *E. coli* and *B. bacteriovorus.* Purple: DAPI, green: msfGFP, orange: tdTomato. Scale bar: 1 µm. **B.** Representative images of *B. bacteriovorus* (strain GL673) expressing *dnaN-mcherry* from the native *dnaN* locus, showing replication initiation (DnaN focus) 50 minutes after co-incubating *E. coli* and *B. bacteriovorus*. Scale bar: 1 µm. **C.** Cell counts expressed in percentages of the samples prepared for Hi-C before and after gradient separation. Total number of cells in each sample before gradient separation: T20: 388, T50: 507, T90: 1206, T120: 140. Total number of cells in each sample after gradient separation: T20: 248, T50: 222, T90: 209, T120: 166, T260: 219.

**Figure S2. A.** Proportion of RNA-seq reads mapped to *E. coli*, *B. bacteriovorus*, or that could not be mapped, shown for all three replicate samples. **B.** Correlation (R-squared) between each sample. **C.** PCA plot of the individual samples shows clustering of time point replicates. Replicates for T0 (attack phase cells isolated after overnight prey-predator culture) and T360 (attack phase cells isolated right after prey exit in a synchronized predation cycle) cluster together.

**Figure S3. A.** Percentage of reads after Hi-C that were mapped to *B. bacteriovorus*, *E. coli,* or that could not be mapped. **B.** Pearson correlation and hierarchical clustering of each Hi-C sample to each other show similarity between attack phase samples, early growth phase samples, and late growth phase samples.

**Figure S4.** Top**:** Hi-C maps, normalized Hi-C signal (blue), and normalized RNA-seq signal (red) at each sampled time point. Middle**:** Zoomed in maps from –946 to 946 base pairs distance from *ori.* Bottom**:** Zoomed in maps from –150 to 150 base pairs distance from *ori.* Arrows point to chromosomal regions strongly enriched in short-range contacts, near *ori* and a three-quarter position locus.

**Figure S5.** Workflow for determining the N/C ratio and median intensity profiles. Denoised images of bdelloplasts obtained by infecting *E. coli* MG1655 with *B. bacteriovorus* strain GL1462 producing cytoplasmic tdTomato were fed into MiSiC [1]. MicrobeJ [2] was used to measure the cell dimensions and BactMAP [3] was used to combine the information from MicrobeJ and the raw images. R was used for further analysis.

**Figure S6. A.** Time-course imaging of the predatory cycle of *B. bacteriovorus* strain GL2870 producing msfGFP-tagged HUα (single copy, native locus expression) and constitutive cytoplasmic tdTomato, showing representative cells for each time point. Cell outlines were drawn manually. Scale bar: 1 µm. **B.** N/C ratio determined using HUα-msfGFP as a proxy for the nucleoid. **C.** Visualization of the median distribution of the HUα-msfGFP signal at each time point. Scale bar: 1 µm. Parameters were the same as for determining the N/C ratio and the median distribution difference of DAPI vs tdTomato in Figure 2.

**Figure S7. A.** Correlation between ChIP-seq read counts per 50 base pairs and RNA-seq reads per 50 base pairs. Sample taken at T0 (AP). Black line: linear model. Pearson R = 0.50. **B.** Correlation between ChIP-seq read counts per 50 base pairs and RNA-seq reads per 50 base pairs. Sample taken at T120 (GP). Black line: linear model. Pearson R = 0.26.

**Figure S8.** Titration curves used to determine the MIC of chloramphenicol and rifampicin. MIC was determined as the concentration where the final *B. bacteriovorus* population was halved compared to a population without antibiotic added. $P_{max}$ was normalized to the $P_{max}$ of the samples without antibiotics per biological replicate. Per experiment, each concentration was measured in triplicate. Dot/triangle shapes indicate separate biological replicates.

**Figure S9.** Correlation of the attack-phase proteome dataset of Hocher et al [4] to all RNA-seq sets in our dataset. Average FQ intensity per gene against fragments per kilobase per million mapped fragments (FPKM) per gene. Pearson correlation coefficients for each sample: T0: 0.05, T20: 0.17, T50: 0.23, T90: 0.22, T120: 0.26, T260: 0.45, T360: 0.07.

**Supplementary references**

1. Panigrahi, S., Murat, D., Gall, A.L., Martineau, E., Goldlust, K., Fiche, J.-B., Rombouts, S., Nöllmann, M., Espinosa, L., and Mignot, T. (2021). Misic, a general deep learning-based method for the high-throughput cell segmentation of complex bacterial communities. Elife *10*, e65151. https://doi.org/10.7554/elife.65151.

2. Ducret, A., Quardokus, E.M., and Brun, Y.V. (2016). MicrobeJ, a tool for high throughput bacterial cell detection and quantitative analysis. Nature Microbiology *1*, 16077. https://doi.org/10.1038/nmicrobiol.2016.77.

3. Raaphorst, R. van, Kjos, M., and Veening, J.-W. (2020). BactMAP: An R package for integrating, analyzing and visualizing bacterial microscopy data. Mol. Microbiol. *113*, 297–308. https://doi.org/10.1111/mmi.14417.

4. Hocher, A., Laursen, S.P., Radford, P., Tyson, J., Lambert, C., Stevens, K.M., Montoya, A., Shliaha, P.V., Picardeau, M., Sockett, R.E., et al. (2023). Histones with an unconventional DNA-binding mode in vitro are major chromatin constituents in the bacterium Bdellovibrio bacteriovorus. Nat. Microbiol., 1–14. https://doi.org/10.1038/s41564-023-01492-x.
